# Spatial transcriptomic analysis of Sonic Hedgehog Medulloblastoma identifies that the loss of heterogeneity and promotion of differentiation underlies the response to CDK4/6 inhibition

**DOI:** 10.1101/2023.02.15.528116

**Authors:** Tuan Vo, Brad Balderson, Kahli Jones, Guiyan Ni, Joanna Crawford, Amanda Millar, Elissa Tolson, Matthew Singleton, Onkar Mulay, Shaun Walters, Marija Kojic, Thomas Robertson, Dharmesh D. Bhuva, Melissa J. Davis, Brandon J. Wainwright, Quan Nguyen, Laura A. Genovesi

## Abstract

**Background:** Medulloblastoma (MB) is a malignant tumour of the cerebellum which can be classified into four major subgroups based on gene expression and genomic features. Single cell transcriptome studies have defined the cellular states underlying each MB subgroup, however the spatial organisation of these diverse cell states and how this impacts response to therapy remains to be determined.

**Methods:** Here, we used spatially resolved transcriptomics to define the cellular diversity within a sonic hedgehog (SHH) patient-derived model of MB and identify how cells specific to a transcriptional state or spatial location are pivotal in responses to treatment with the CDK4/6 inhibitor, Palbociclib. We integrated spatial gene expression with histological annotation and single cell gene expression data from MB, developing a analysis strategy to spatially map cell type responses within the hybrid system of human and mouse cells and their interface within an intact brain tumour section.

**Results:** We distinguish neoplastic and non-neoplastic cells within tumours and from the surrounding cerebellar tissue, further refining pathological annotation. We identify a regional response to Palbociclib, with reduced proliferation and induced neuronal differentiation in both treated tumours. Additionally, we resolve at a cellular resolution a distinct tumour interface where the tumour contacts neighbouring mouse brain tissue consisting of abundant astrocytes and microglia and continues to proliferate despite Palbociclib treatment.

**Conclusions:** Our data highlight the power of using spatial transcriptomics to characterise the response of a tumour to a targeted therapy and provide further insights into the molecular and cellular basis underlying the response and resistance to CDK4/6 inhibitors in SHH MB.

## Background

Medulloblastoma (MB) is the most frequent malignant paediatric brain tumour and the leading cause of cancer-related mortality and morbidity in children[63]. Current treatment options consisting of surgical resection, craniospinal irradiation and high-dose chemotherapy have improved survival rates to approximately 70-75% for children with average-risk MB [20, 22, 64]. These treatment options are still ineffective however for a substantial number of patients, such as those diagnosed with high-risk disease and those that go on to relapse, a universally fatal prognosis[34, 67, 69]. Moreover, survivors face significant long-term neurocognitive, endocrine and physical sequalae as a consequence of aggressive treatment protocols. Clearly, there is an urgent need for improved, targeted therapies that minimise these harmful side effects.

Tumour heterogeneity is one of the major challenges limiting the efficacy of effective targeted therapy. Bulk-tumour transcriptomic, methylomic and genomic profiling studies focused on the MB at diagnosis indicate extensive genetic intertumoural heterogeneity. Four consensus subgroups of MB are now recognised, Wingless (WNT), Sonic Hedgehog (SHH), Group 3 (Gp3) and Group 4 (Gp4) [62], with subsequent studies further subdividing these subgroups into a number of subtypes [10, 61, 74]. The stratification of patients on the basis of distinct molecular features defining MB subgroups has a profound impact on their clinical outcome (as reviewed by [68]) resulting in subgroup-specific profiles bring incorporated into risk-adapted therapy stratification in current biomarker-driven clinical trials of upfront therapies (SJMB12, NCT01878617, NCT02066220)[63]. However, not every genetic lesion will necessarily be present in every cell of a tumour, with refractory or relapsed disease likely arising due to the failure of therapy to eradicate all cell types. Indeed, both molecular subgroup and novel subtypes remain stable in the majority of relapsed disease[69], and relapsed MB shown to be frequently driven by a dominant clone arising through clonal selection of a pre-existing genotype present at diagnosis [34]. A greater understanding of therapy-resistant cells at diagnosis and their evolution throughout tumour progression and therapy is urgently needed.

Cellular heterogeneity is a key factor in cancer progression, resistance and relapse, in that individual tumour cells having different responses to treatment allowing certain cells to continue to expand under therapeutic selection pressure[74]. Thus, characterising cellular diversity is necessary to gain a deeper understanding of tumour progression and therapeutic response. Early evidence for intratumoural heterogeneity in MB came from immunohistochemical (IHC) studies, with highly heterogeneous patterns of key biomarkers among cells of the same tumour[17, 19, 34]. Histological studies support this heterogeneity, with the distinction of histopathological MB subtypes often problematic due to complex mixtures of subtypes within the same tumour [46].

Recent single cell RNA sequencing (scRNA-seq) studies have begun to resolve the complex cell type composition of MB [36, 70, 77, 84], identifying immune cell infiltrates [70] and subgroup-specific neoplastic subpopulations defined by cycling, progenitor and differentiated neuronal programs [36, 70, 77]. Consensus biomarkers of MB subgroups were restricted to discrete subpopulations of cells that are only present in certain MB subtypes of a particular subgroup [70], therefore not accurately representing all subtypes within a certain MB subgroup. While these scRNA-seq studies have been transformative for resolving the cellular states within subgroups of MB, tissue dissociation leads to the loss of information regarding the spatial organisation of cells within the tumour microenvironment (TME). Spatial variability within the TME has been shown to substantially impact tumour cell invasiveness [39], therapeutic response [6] and clinical outcomes [40, 73]. Therefore, understanding the complex relationship between MB cell types and their relative spatial features within the TME is crucial for developing a greater understanding of functionality of MB cell types and their prognostic value.

Spatial Transcriptomics sequencing (ST-seq) has recently emerged as a technology to address the limitations of both bulk and scRNA-seq, providing whole transcriptome analysis across intact tissue sections without the need to dissociate cells from their *in situ* localisation within the tissue. The advent of ST-seq technologies offers an unprecedented opportunity to better characterise complex interactions within the tumour microenvironment at a high resolution and therefore identify intratumoural spatial neighbourhoods of both biological and therapeutic relevance [25].

Palbociclib (IBRANCE, Pfizer, Inc) is a selective CDK4/6 inhibitor that functions to block Retinoblastoma hyperphosphorylation and the E2F-mediated transcriptional program, arresting cells in the G1 phase of the cell cycle [21]. Previously, we demonstrated that Palbociclib treatment results in a highly significant survival advantage to mice bearing Med-1712FH (SHH) patient-derived orthotopic xenograft (PDOX) MB [15], although 80% of the tumours relapse upon the cessation of treatment. Using a SHH PDOX model of MB, here we optimised a ST-seq approach to define the transcriptional state of cells and their spatial locations within the TME and compared untreated tumours to tumours undergoing treatment with Palbociclib. Integrating spatial gene expression with histological data, we develop an analysis strategy to investigate the hybrid system of mouse and human cells and their interface within the cerebellum. With spatial measurements of thousands of genes, we have performed unbiased genome-wide and pathway-level analyses and show that Palbociclib induces a differentiated neuronal program in the majority of the tumour but that an interface of mixed tumour cells continue to display a proliferative transcriptional state at the tumour/microenvironment boundary. Overall, this study demonstrates the power of a ST-seq approach in quantifying and spatially mapping MB spatially heterogeneous responses to therapies within the appropriate microenvironment. In turn, this provides a nuanced understanding of the inter-relationship between tumour heterogeneity, therapeutic response and the TME.

## Methods

### Mice

Seven to nine week old male NOD.Cg-*Prkdc*^scid^ *Il2rg^tm1Wjl^/SzJ* (NSG) mice (Jackson Laboratory, Bar Harbor, ME) were maintained in a barrier facility on a 12-hour light/dark cycle with food and water provided ad libitum, in accordance with the NIH Guide for the Care and Use of Experimental Animals. All experiments were performed with approval from The University of Queensland Molecular Biosciences animal ethics committee (IMB/386/18).

### Medulloblastoma PDOX mouse model

The Medulloblastoma PDOX mouse model Med-1712FH used in this study was generated in the Olson laboratory (Fred Hutchinson Cancer Research Center, Seattle) using paediatric patient tumour tissue obtained from Seattle Children’s Hospital with approval from the Institutional Review Board. This model is publicly available from https://research.fredhutch.org/olson/en/btrl.html, with details of this model published [7]. Med-1712FH tissue was derived from a 4.9–year old patient and classified as a SHH-MB, characterised by desmoplastic/ nodular morphology. The PDOX line was generated by implanting the tumour cells in the cerebellum of immunocompromised NSG mice within hours of surgical removal from the patient and propagating them from mouse to mouse exclusively without in vitro passaging as previously described[7]. Xenografted tumours were subjected to genomic analysis and compared to the primary tumour from which they originated[7].

To generate orthotopic xenografts, NSG mice were anesthetized and a small incision made in the skin to expose the skull. A handheld 0.7 millimetre microdrill was used to create a hole in the calvarium above the right cerebellar hemisphere, two mm lateral (right) to the sagittal suture and two mm posterior of the lambdoid suture. Tumour tissue from symptomatic donor mice was dissociated in serum-free Dulbecco’s Modified Eagle’s Medium (DMEM) using a 21G needle. The cells were filtered, centrifuged, and resuspended in DMEM at a concentration of 50,000 cells/ul. Two microlitres of the cell suspension (100,000 cells) was injected into the brain parenchyma approximately two millimetres under the dura. SurgiFoam was inserted into the burr hole site to mitigate leakage and the incision was closed with surgical staples.

### In vivo systemic chemotherapy

Brain tumour growth was confirmed in each mouse by the onset of symptoms and randomly assigned into treatment groups. Mice were enrolled in the study at a score of four and were observed with weights recorded daily. Palbociclib hydrochloride (Pfizer) was used for all experiments in this study. Palbociclib was dissolved in 50 mmol/L sodium lactate, pH 4 and administered orally daily at 100mg/kg for a total of seven days before being collected for either Visium Spatial Gene Expression or immunofluorescence. This timepoint corresponds to early treatment where we first see visible signs of drug efficacy (data not shown) therefore correlating to potential changes to intratumoural heterogeneity at this point. For all untreated tumour-bearing mice, animals were euthanatized prior to the end of the study if mice demonstrated signs of tumour-related morbidity or lost more than 20% body weight loss.

### Sample embedding and quality control

Brain samples were embedded in the optimal cutting temperature compound (OCT) immediately following the dissection. OCT-covered brain specimens were frozen by dipping into liquid nitrogen chilled isopentane solution. OCT-embedded tissue blocks were cryosectioned at eight micron (μm) thickness. Three to four sections were stored at -80°C prior to RNA extraction using the QIAGEN RNeasy micro kit (#74004). RNA yield was measured by Qubit RNA HS assay kit (#Q32852), with quality assessed using an Agilent 2100 Bioanalyzer with RNA 6000 Pico assay (#5067-1513). The RNA integrity number (RIN) of all samples was approximately nine. The remaining sections were placed onto SuperFrost Plus (#J1800AMNZ) slides for the assessment of sample morphology and optimisation of H&E staining (Fig. S1). Tissue fixation and staining were as described in the ‘Methanol Fixation, H&E Staining and Imaging’ Visium protocol (10X Genomics; #CG000160), with variations being haematoxylin staining for five minutes (min), bluing for one min and eosin staining for two mins. Stained sections were imaged on a Zeiss AxioScan F1 Fluorescent Slide Scanner at 20X magnification.

### Visium Spatial Tissue Optimisation

Sample-specific tissue permeabilisation optimisation was performed following the Visium Spatial Tissue Optimisation User Guide Rev B (10x Genomics, #CG000238). Tissue optimisation aims to maintain the balance between transcript capture efficiency and RNA diffusion. This process is especially critical for tumour tissue, comprising regions of densely packed nuclei which are difficult to permeabilise. Additionally, human cancer tissue and the host mouse brain have varying permeabilisation efficiencies. To address this, we compared sections of varying thickness in addition to the trialling several permeabilisation times. Ultimately, eight micron sections were determined as the optimal thickness for both tissue types and selected for downstream analysis. Sequential cryosectioned frozen tissue sections were placed inside each of eight capture areas on a Visium Tissue Optimisation Slide (#3000394), fixed in pre-chilled methanol at -20°C for 30 mins, stained in Mayer’s hematoxylin (Dako) for five mins and eosin (Sigma) for two mins. Imaging was performed on a Zeiss Axio Scan Z1 slide scanner. Tissue in the eight arrays was then permeabilised over a range of time points (one to 70 mins) to allow cellular RNA to hybridise directly to the oligo-dT nucleotides printed on the slide (Fig. S1). The captured RNA was reverse transcribed, incorporating fluorescently labelled nucleotides to generate fluorescently labelled complementary DNA (cDNA). The fluorescent oligonucleotide-bound cDNA was visualised on a Leica DMi8 inverted widefield microscope. The H&E and fluorescent images were then assessed to select an ideal permeabilisation time that generated the brightest and most distinct signal, correlating to morphological features of the tissue (Fig. S1). Optimised conditions of eight microns thickness with eight mins permeabilisation were then used for Visium Gene Expression.

### Visium Spatial Gene Expression Library Preparation

Sequencing libraries were prepared according to the Visium Spatial Gene Expression User Guide Rev C (10X Genomics, #CG000239). Visium Gene Expression Slides (#2000233) contain four capture arrays with oligonucleotides for RNA capture, spatially distinguished by approximately 5000 uniquely barcoded ‘spots’ (Fig. S1). These barcoded oligonucleotides also contain Unique Molecular Identifiers (UMIs) to allow tracing of single mRNA transcripts per spot. Eight micron MB PDOX tissues were sectioned onto a Visium Gene Expression Slide and the tissues were permeabilised for eight mins to allow release of RNA onto the slide. Following reverse transcription, amplification of cDNA was performed for 17 cycles and final indexed samples were amplified for 12 cycles. All libraries were loaded at 1.8pM onto a NextSeq500 (Illumina) and sequenced using a high output reagent kit (Illumina) at the Institute for Molecular Bioscience Sequencing Facility. Sequencing was performed using the following protocol: Read1 - 28bp, Index1 - 10bp, Index2 - 10bp, Read2 - 120bp.

### Read Mapping

Illumina sequencing base call data (BCL) was converted to FASTQ files using bcl2fastq/2.17. The resulting FASTQ reads were trimmed to remove poly-A sequences on the 3’ end and any remaining of the template switch oligo (TSO) sequences on the 5’ end of the Read 1 files by using cutadapt/1.8.3.^32^ The cleaned FASTQ files were then used for mapping by SpaceRanger V2.0 (10x Genomics), based on STAR for splicing-aware alignment. A hybrid genome refence sequence was created by combining the human reference genome (GRCh38-3.0.0) and mouse reference genome (GRCm38 - mm10-3.0.0), using SpaceRanger 1.2.2. Mapping was performed with SpaceRanger 1.2.2 (STAR 2.7.2a) and the sequenced data was then mapped to the spatial coordinates in the H&E image based on spatial barcode information. Only spots within the tissue region were included for downstream analysis, with reads mapping to spots outside the tissue region discarded. Reads mapping outside the tissue region are often due to ambient RNA from the tissue sectioning procedure prior to methanol fixation on the slide. By default, the all reads outside the tissue are removed in the SpaceRanger pipeline and the resulting filtered matrix output from SpaceRanger is used for ongoing analysis.

### Assessing the quality of the spatial data by comparing to mouse brain anatomy

To evaluate data quality, we investigated whether we could identify known cerebellum structures using unbiased clustering of gene expression data, using stLearn v0.3.1 [65]. We filtered genes with fewer than 50 counts, and log transformed the data. Spatial morphological gene expression (SME) normalisation was then applied to improve individual spot expression quality; whereby neighbouring spots with similar gene expression and histological features were used to correct for noisy gene expression measurements[65].Louvain clustering (resolution = 0.75) was performed on a neighbourhood graph constructed using 50 nearest neighbours based on the top 50 principal components calculated after SME normalisation as per the standard SME clustering workflow in stLearn. To identify gene markers distinguishing a cluster from the remaining cells within a sample, all genes in each cluster were ranked using a one-versus-rest approach with Wilcoxon rank-sum tests in scanpy v1.8.2 [80] (Table S1). For each cluster, the top 10 genes that were differentially expressed compared to remaining clusters (excluding mitochondrial genes) were then used for Giotto enrichment analysis (detailed below) to assess the specificity of the gene markers for the cerebellum structures of interest across each sample (Fig. S2).

### Spatially stratifying spots by species and tissue regions

To label each spot according to the respective species, we first filtered a total of 292 low quality spots which had fewer than 200 genes detected, removing approximately two percent of all spots (Fig. S3). We then merged all spots across samples into a single *Seurat* v4.0.1 object, and applied *sctransform* normalisation [31] across genes and samples. For each spot, two ‘species scores’ were calculated by summing the normalised and unscaled gene expression for human or mouse genes separately to avoid the effect of outlier over-abundant genes. The scores were used to classify spots as either ‘human’, ‘mouse’, or ‘mix’; with the latter representing spots containing cells from both species (Fig. S4a) as described below.

For each sample, the species labels were determined as shown in equations 1-3:

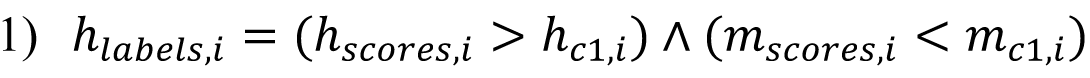

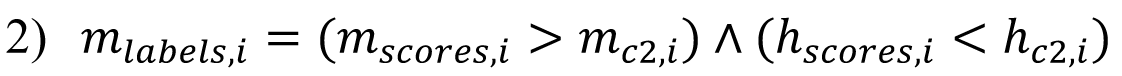

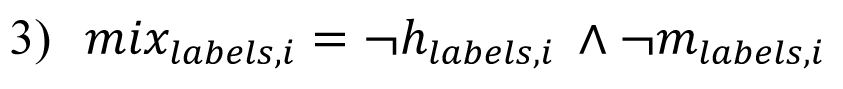

Where #x1D456; indicates the sample, ℎ_!"#$!%,’_ indicates the human labelled spots, ℎ_%()*$%,’_ represent the human scores across the spots, #x1D45A;_!"#$!%,’_ represent the mouse labelled spots, #x1D45A;_%()*$%,’_ represent the mouse scores across spots, #x1D45A;#x1D456;#x1D465;_!"#$!%,’_ represent the mix labelled spots. . ℎ_(+,’_, ℎ_(,,’_, #x1D45A;_(+,’_, #x1D45A;_(,,’_ are cutoffs to classify spots, as determined by score distribution shown in (Fig. S4) and values listed in (Table S2).

Based on the distribution of the human/mouse scores in the scatter plots and correspondence between species classification and tissue histology, cut-offs were manually chosen for each sample (Figure 1, Fig. S4a). For sample Control C, a small number of spots outside the tumour region had low levels of human gene expression (Fig. S4e) and no clear tumour histology was identified on pathological inspection (Fig. S4e), so to prevent mislabelling these outlier spots, we subsequently labelled these as ‘mouse’ spots.

**Figure 1.**
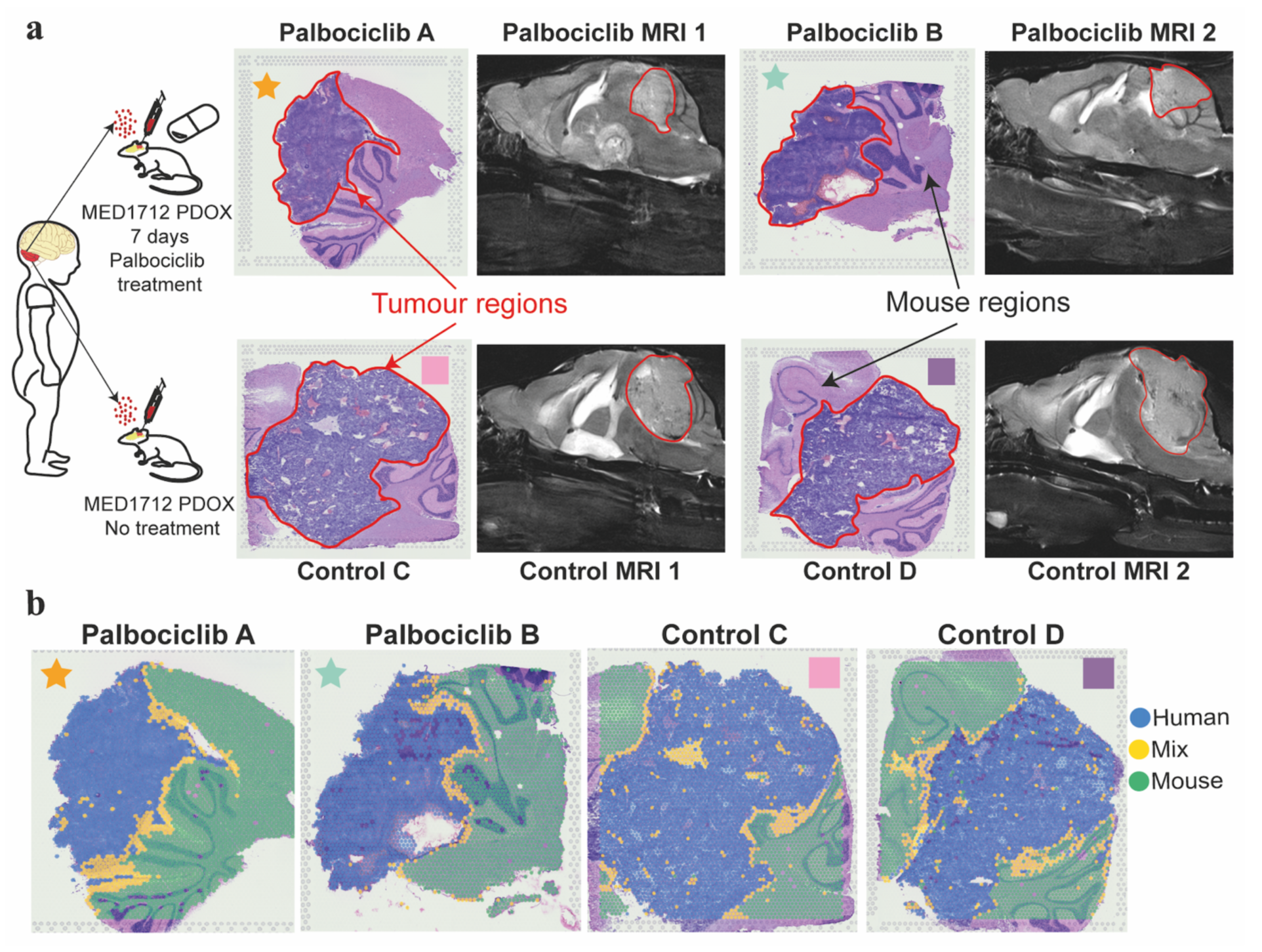
Visium spatial transcriptomics detects spatial human tumour and mouse transcripts within Medulloblastoma PDOX undissociated tissue sections. (a) Illustration of experimental design. Sonic hedgehog PDOX medulloblastoma (Med-1712FH) mouse models were generated by seeding tumour cells in mouse cerebellum. Mice received a seven day course of Palbociclib and compared to untreated control tumours. H&E stained images display regions of human tumour and mouse brain in Palbociclib-treated mice (demarcated ‘Palbociclib A’ and ‘Palbociclib B’) and controls (demarcated ‘Control C’ and ‘Control D’). Contrast-enhanced MRI shows the tumour location in PDOX mouse brain (outlined in red). Palbociclib-treated and control sections are indicated with stars and squares, respectively. (b) The spatial context of the human, mix, and mouse spots. Spots are classified as human (green), mix (yellow) or mouse (blue) based on whether they predominantly express one species’ genes or a combination of both. PDOX Patient-derived orthotopic xenograft, MRI magnetic resonance imaging.

### Bioinformatics analysis of gene expression markers

The voom pipeline [48] from limma v3.48.1 was used to identify differentially expressed genes between treated and untreated conditions using “pseudobulked” samples of each species. For pseudobulking [53] analysis, we summed UMI counts for each gene across all spots belonging to a particular species and sample. We then subsetted to genes belonging to the relevant species; only human genes were considered for human spots, only mouse genes were considered for mouse spots, and both human and mouse genes were considered for mixed spots (Fig. S4b). For the mixed spots, differential expression (DE) analysis was performed separately for human and mouse genes.

The data were transformed to log(counts per million(CPM)+0.5), hereby denoted as log-cpm, using edgeR v3.34.0 to perform a density based filtering of lowly expressed genes (Fig. S4b). Removal of lowly expressed genes was then performed separately for pseudobulked counts for each species using the same workflow (Fig. S4c). This included filtering of genes with no detectable expression across all spots in all samples and removal of genes with a maximal level of expression below a particular threshold. Minimum thresholds were chosen using density plots of the gene expression distribution as a diagnostic plot to remove lowly expressed genes. For human, mouse, mix-human genes, and mix-mouse genes; the minimum log-cpm thresholds used were two, two, three, and three (due to the higher distribution of the first mode), respectively. The higher cutoffs for the mix-human and mix-mouse was due to the shifted distribution of log-cpm values (Fig. S4b); whereby the peak representing the lowly expressed genes was at a higher log-cpm value than in the human and mouse data. After filtering lowly expressed genes, a total number of 13069, 13700, and 21182 (11308 mouse, 9874 human) genes remained for DE testing in the human, mouse, and mixed pseudobulked data, respectively (Table S3-5).

After gene filtering, trimmed-mean-squares (TMM) [71] was applied to determine robust size factors for adjusting log-cpm values. TMM trims genes based on log-FC between samples before size factor estimation, a calculation which results in a lower accuracy if lowly expressed genes are not removed beforehand. The gene expression distributions of the samples were clearly equivalent after the above steps were taken, indicating the data was appropriately normalised (Fig. S4d). Relative Log Expression (RLE)[23] plots were generated to assess the presence of unwanted variation in the data. Median RLEs across spots were mostly zero suggesting that unwanted sources of variation were minimal and unlikely to skew downstream statistical analysis (Fig. S4g). For each of the pseudobulked species profiles, we then performed mean-variance correction using voom with treatment contrasted against the control to ensure downstream DE analysis with limma was not affected by mean gene expression [53]. The TREAT criterion [56] was applied to identify genes with absolute logFC significantly greater than 0.15 (Fig. S4f). Genes were considered differentially expressed if they had a false discovery rate corrected p-value < 0.05.

### Gene Set Enrichment Analysis (GSEA)

GSEA was performed on the human DE genes (comparing the human region between treated and control samples) against the Molecular Signature Database (MSigDB) Hallmark and Biological Process Gene Ontology gene sets. DE genes were ranked from most upregulated to most downregulated genes using t-values, which estimate the magnitude of differences between two means relative to variation when comparing treated and control samples. GSEA was performed using the GSEA function from clusterProfiler v4.0.0 with a minimum gene set size of 20, using fgsea method[81], 10000 permutations and a 0.01 adjusted p-value cutoff. The full list of enriched terms are provided in Table S6.

### Estimating geneset enrichment scores per spot

To understand the spatial activity of gene sets, we performed Giotto spot enrichment analysis using DE genes which overlapped with the indicated gene sets (see Results). We utilised the sctransform normalised data subsetted to the relevant species genes as input for the enrichment analysis. The method used was Parametric Analysis of Gene Expression (PAGE). Briefly, PAGE calculates per-spot enrichment scores by first determining the fold-change of each gene within a spot relative to the mean expression of that gene across all spots [44]. For an inputted gene set, the gene set enrichment score (#x1D438;) for a given spot is calculated based on the expression fold-changes of genes within the gene set #x1D446;. relative to the expression of these genes in all the remaining spots. The fold-changes are normalised by the mean fold-change (#x1D707;) of all (#x1D446;.) genes in the spot, the standard deviation of these fold-changes (#x1D6FF;) in the spot, and the number of genes within the gene set (#x1D45A;) using equation 4.

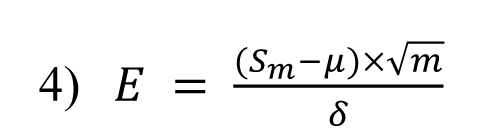

A spot with a high E value suggests the gene set is more active in that spot compared to in the remaining spots. The function PAGE Enrich in Giotto version 1.0.3[18] was used to calculate this enrichment score per spot using expression_values=’normalised’ as input. The enrichment scores reflect how much higher or lower a spot expresses a gene set relatively to all other spots within the tissue.

### Automated spot annotation by dominant cell type

SingleR v1.6.1 [2] was used to map a dominant reference cell type to each spot based on Spearman’s correlation coefficient and nearest label classification. For human/mix spots, only human genes were used to map against annotated human reference scRNA-seq data (Fig. 5g, Fig. S7a). Similarly, for annotating mouse/mix spots, only mouse genes were used to map against annotated mouse reference scRNA-seq data (Fig. 2d, Fig. S5-6). For both the human and mouse annotations, each sample was analysed independently. The data was processed as described above, removing genes with expression in fewer than three spots, followed by log(CPM+1) transformation. For spot annotation analysis, we utilised spatial and tissue morphology information to adjust for gene expression values in the way that spots in close proximity and with similar morphology (e.g. number of nuclei and spatial distribution of nuclei). This way, we could identify clusters of spots that are similar in spatial location, morphology and gene expression. To achieve this, the Spatial Morphological gene Expression (SME) adjustment method in stlearn v0.3.1 [65] was applied. Only shared genes detected in both spatial samples and reference scRNA-seq data were used for SingleR cell mapping.

**Figure 2:**
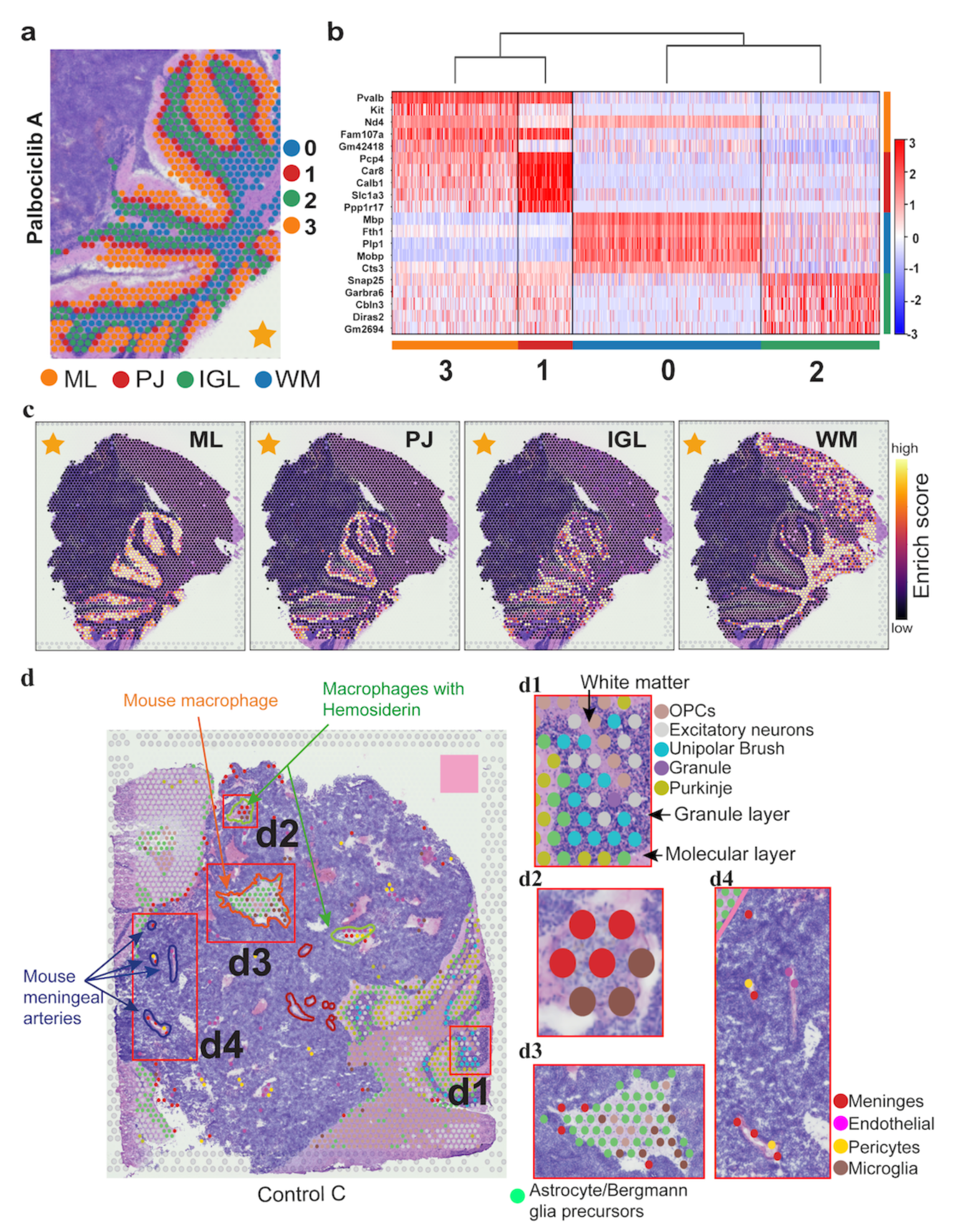
High-resolution functional structures of mouse PDOX brain sections can be identified using unsupervised clustering and automated cell type identification based on spatial gene expression. (a) Unsupervised clustering of mouse spots from cerebellum regions in PDOX sections based on gene expression. By this data-driven approach, each cluster precisely corresponded to a known anatomical layers of mouse cerebellum. (b) Heat map of cluster marker genes depicts the top five differentially expressed genes in each cluster. (c) Spot gene enrichment scores of top ten most differentially expressed genes in each cluster, consistent with clustering results shown in a, but with additional information on gene expression heterogeneity within each cluster. (d) High-resolution cell types identified by reference-based annotation (see Methods) were concordant with and expanded upon pathological annotation of the H&E sections. Independent annotations by a brain pathologist for the H&E tissue image are shown as close contours overlaid with the mouse/mix spot cell types. Sample control C is shown as a representative. (d1-d4) Enlarged regions of interest from d (red boxes) showing mouse cell types consistent with distinct histological features, but with higher resolution reflecting important heterogeneity. (d1) Dominant cell types indicated at each layer are as expected, with oligodendrocyte precursor cells in the white matter (inner-layer), granule and unipolar brush cells in the granule layer (dark, middle layer), and Purkinje cells in the molecular layer (outer layer). (d2) ‘Macrophage with Hemosiderin’ pathological feature contained exclusively meninges and microglia. (d3) ‘Mouse macrophage/lymphocytes’ area was predominantly astrocytes, microglia, and meninges. (d4) ‘Mouse meningeal arteries’ corresponded to vasculature cell types; meninges, endothelial cells, pericytes, and microglia. IGL Internal Granule Layer, ML Molecular layer, OPC Oligodendrocyte Precursor Cells, PJ Purkinje, WM, White Matter.

In the case of human spot annotation, the reference scRNA-seq data was 13-week old male human foetal brain from the Human Cell Landscape database (Fig. S7a; Foetal brain 3, available at http://bis.zju.edu.cn/HCL/dpline.html?tissue=Brain). The reference count data was normalised using SCTransform (Seurat v4.0.1) and log-transformed before the SingleR run. The transformed gene expression of the reference single cells within each cell type annotation was pooled and averaged to create 3 reference gene expression profiles per cell type using the ‘aggregateReference’ function in SingleR with ncenters=3. Genes were then selected for subsequent auto-annotation using the aggregated reference profiles and the ‘trainSingleR’ function with 350 top DE genes (identified by SingleR) sorted by log foldchange (de.method=’classic’). Using the selected genes and the aggregated cell type reference profiles, spots within each sample were then matched to all cell types in the reference. Spots were given the annotation of a cell type for which they had the highest Spearman’s correlation coefficient. This was performed via the ‘classifySingleR’ function.

For the mouse spot annotation, the reference scRNA-seq data was developing mouse cerebellum data from Vladiou, *et al.* (2019) sourced directly from the Gene Expression Omnibus (GEO) (GSE118068). The murine cell types in this dataset were provided by the original publication (Fig. S7b). Genes expressed in less than ten cells were removed from the reference data and the data was transformed to log(CPM+1) using scanpy v1.8.1. In order to capture the most relevant cell types for our adult mouse tissue we subsetted the reference data to only include cell types present at the earliest available time point, i.e. postnatal day 7 (Fig. S6b). Only shared mouse genes across the mouse/mix spots per sample and the reference scRNA-seq were used. For each cell type, the mouse scRNA-seq data was equivalently aggregated as described for the human tissue annotation. Genes were subsequently selected for the pattern matching using the ‘trainSingleR’ method, with parameters de.n=200 and de.method=’t’. Parameters were chosen so that known cell types mapped to the correct histological structures of the cerebellum as a diagnostic for accurate cell type detection (Fig. 2d, Fig. S5).

#### Cell type enrichment analysis at tumour-mouse brain interface

Border regions defined as “mixed” spots at the human-mouse interface region were used for cell type enrichment analysis at tumour-mouse brain interface (Fig. S6b,c,d,e). For each sample, a contingency table was constructed by counting the number of ‘Astrocyte/Bergmann glia’ and non-‘Astrocyte Bergmann glia’ spots in the border region and in the remaining mouse tissue (Table S7), used as input for Fisher’s exact test for a cell type in the border versus in the tumour region (scipy v1.6.2, alternative=’greater’) (Table S7).

### Heterogeneity analysis

To study heterogeneity among human spots, we quantitatively calculated four different scores including Connectivity index, Shannon entropy, Simpson index, and cluster modularity. For initiation, we used the graph-based Louvain community detection method to identify changes in cell clustering results between 23 different clustering resolution, with a smaller resolution resulting in smaller cell communities detected and more clusters. This means that the number of clusters and spots allocation alters among different clusters between different resolutions, reflecting the level of heterogeneity in the gene expression data among the spots. For each resolution, the membership of each cell in each cluster is recorded, with the split of two cells belonging to one cluster in a resolution into two clusters in the next resolution summarised and visualised as arrows connecting clusters. We then calculated the connectivity index (CI) as a measure of heterogeneity due to separating or grouping spots that are similar defined as nearest neighbours, reflecting degree that similar spots are in the same cluster. A higher CI suggests high heterogeneity [33].

To further measure heterogeneity quantitatively, we calculated two information-theory measures, namely Shannon entropy and Simpson index of gene expression across human spot clusters in each of the clustering resolutions [43]. Briefly, the two scoring methods calculated the probability that any two spatial spots belong to one homogenous cluster. Shannon entropy represents the uncertainty of spots grouped into one cluster, with high Shannon entropy indicating high heterogeneity. In contrast, Simpson index computes the probability that two randomly selected spots belong to the same cluster, and thus a low Simpson index indicates high heterogeneity. We also calculated cluster modularity, reflecting purity (homogeneity) of spots within clusters [43, 54].

## Multiplex single molecule RNA-*in situ* hybridization (smRNA-FISH/RNAscope)

OCT-embedded tissue blocks of Med-1712FH PDOX were sectioned on a crystat (Leica) at eight µm and mounted on Superfrost Plus slides (ThermoFisher). Cryosections were stored at −80°C until further use. 8-plex single molecule RNA-FISH (smRNA-FISH) was performed RNAScope Hiplex assay (ACDbio/biotechne) as outlined in the RNAScope HiPlex Assay User Manual (324100-UM). mRNA target probes for transcripts of interest were designed by ACDbio probe design team and performed using RNAscope HiPlex assay (ACDbio Cat. No. 324110): *Interleukin 4* (*Il4)* (1143601-T3), *Interleukin 4 Receptor* (*Il4ra)* (1085871-T4),

CD68 Molecule (Cd68) (1143581-T5), Allograft Inflammatory Factor 1 (Aif1) (1143591-T6), Insulin Like Growth Factor 1 (Igf1) (1143641-T7), S100 Calcium Binding Protein B (S100b) (1148311-T10), Marker Of Proliferation Ki-67 (MKI67) (548881-T11), Glial Fibrillary Acidic Protein - Gfap (313211-T12). A set of negative control probes (Cat no. 324341) were used for staining of negative control slide to assess for non-specific binding of amplifiers used in fluorescent labelling. Briefly, sections were fixed in freshly made 4% paraformaldehyde for one hour, washed two times with PBS and dehydrated in Ethanol. Sections were then digested with Protease IV (ACDbio, Cat. No 322336) for 30 min at room temperature and then incubated with targeted-probe mix and amplifiers according to the manufacturers instructions. Sections were stained with DAPI for 30 seconds and mounted in Prolong Gold Antifade (ThermoFisher). Up to four transcripts were labelled per imaging round by AF488, AF568, AF647, AF751 fluorescent dyes. Between imaging rounds, coverslips were removed, and fluorophores of previous imaging rounds were cleaved to enable consecutive rounds of imaging, with each round containing probes for a new set of transcripts.

The images were captured by Zeiss LSM900 with an appropriate adjustment of fluorescence intensity. We used DAPI as reference signal imaging, also to selecte region of interest. A 10% overlap of tiles were set up to ensure accurate stitching. Lastly, the images processed tiles stitching and adjusting contrast/brightness by using by ZEN software (version 3.2). The signals from different round were merged by using HiPlex v2 - Image Registration Software.

### Immunofluorescence analysis of tumours

For immunofluorescence analysis, tumours were preserved by paraformaldehyde fixation and processed for immunoanalysis. Antibody markers were analysed on seven micron, paraffin-embedded sections via standard immunofluorescence techniques using the following antibodies: marker of proliferation (Ki67) (Abcam ab15580, 10ug/ml), Glial Fibrillary Acidic Protein (1:100, Chemicon MAB360), Stathmin 4 (1:50, Proteintech 12027-1-AP) and Tubulin

Beta 3 Class III (1:200, Abcam ab78078). High temperature unmasking was performed with pretreated pH6.0 citrate buffer (Vector Labs) for 5 min, and mouse-on-mouse blocking reagent was used to block non-specific binding of mouse primary antibodies (Vector Labs). Images were captured using Zeiss LSM 710 upright confocal microscope as Z-stacks and presented as the sum of the Z-projection. The images were further exported for processing and analysis in ImageJ[72]. Cell number and mean pixel intensity quantifications were performed per field of view (FoV; scale bars specified in the figure) and three FoV were analyzed per tumour in triplicate for each group. The number of Ki67^+^ cells and a total number of cells (DAPI) were determined manually using ImageJ Cell Counter plugin. TUBB3 and STMN4 expression was quantified as mean pixel intensity per FoV. The graphs were prepared in Prism v9.0.1 (GraphPad) software. The number of replicates, corresponding statistical tests, and statistically significant differences are indicated in figure legends.

### Immunoblot analysis of tumours

Whole cell extracts were generated by lysing cells in radioimmunoprecipitation (RIPA) buffer (1% IGEPAL, 150mM NaCl, 50mM Tris (pH 8), 0.5% Sodium deoxycholate, 0.1% SDS) with protease and phosphatase inhibitors (Cell Signalling Technology, 5872) and 1uL/mL Benzonase® nuclease (Millipore, E1014). Total protein concentrations were determined using the BCA kit (Pierce, 23225) and transferred to a polyvinylidene difluoride (PVDF) membrane using a transfer apparatus according to manufacturer’s protocol (Invitrogen, B1000). After incubation with 5% non-fat milk in TBS-T (20mM Tris, 0.15M NaCl, 0.1% Tween-20, pH 7.6) for 60 min, the membrane washed once with TBST and then incubated with an anti-beta III tubulin (Abcam, ab78078, 1:20,000), anti-Stathmin 4 (United Bioresearch, 12027-1-AP, 1:1000) and anti-Actin (Sigma-Aldrich, A2066, 1:1000) primary antibody for 16 hours at 4°C. Membranes were washed three times for five min and incubated with a horseradish peroxidase (HRP)-conjugated, donkey anti-mouse (Abcam, ab6820, 1:2500) or donkey anti-rabbit

(Abcam, ab16283, 1:2500) secondary antibody for one hour at room temperature. Membranes were then exposed to SuperSignal West Pico PLUS Chemiluminescent substrate (Thermo Scientific, 34580) and imaged on the BioRad ChemiDoc MP Imager. For re-probing of membranes, HRP signals were quenched by incubating membranes in 16.67mM Sodium azide for approximately 14 hours at room temperature with signal quenching confirmed by re-imaging the membrane.

## Results

### Spatially resolved transcriptomics maps a mixed human mouse interface in SHH patient-derived MB

To investigate the spatial organisation of cells in SHH patient-derived MB, we profiled spatial gene expression in Med-1712FH (SHH) PDOX tissue sections using the capture-probe based ST-seq (10X Genomics Visium platform). We employed a PDOX SHH model of MB as this represents a well characterised, accurate model of SHH MB and we have extensively characterised its overall response to Palbociclib. PDOX MB have been shown to maintain the characteristics of the primary human tumours from which they were derived from in terms of histology, immunohistochemistry, gene expression, DNA methylation, copy number and mutational profiles [7]. To monitor how intratumoural heterogeneity changes in response to therapeutic selection, we generated spatial molecular maps combining both imaging and sequencing data from Med-1712FH (SHH) PDOX and compared to these Med-1712FH (SHH) PDOX obtained from mice treated with the CDK4/6 inhibitor, Palbociclib (Fig. 1a).

Our ST-seq dataset contained transcriptomes for 14743 barcoded array spots across the four samples, encompassing a total of 64591 unique human and mouse genes. Each array spot is 55µm in diameter and contains approximately one to nine cells. We applied an automated classification system, based on the number of reads mapped to the human, or to the mouse component of the hybrid reference genome to define a spot as a human only, mouse only or a mixture of both human and mouse cells (see Methods). The datasets represented a comprehensive transcriptome-wide profiling of all common human and mouse genes in a hybrid in *vivo* system. We detected a high number of genes across the tissue, with medians of genes per spot at 1263 -1435 (Fig. S3a, d-h). Mapping of sequencing reads to reference genomes and spatial location across the tissue section clearly demarcated human and mouse tissue regions, strongly recapitulating species origin of the tissue on the basis of histology (Fig. S3b-c). We next devised a quantitative approach to define species-origin of each spatial spot, based on the levels of reads aligned to respective species within the spot. The classification of the majority of spots again closely mirrored the species origin as determined from histology, with the majority of the PDOX labelled as human and the surrounding mouse tissue as mouse (Figure 1b). The high concordance in molecularly defined tissue regions with the distinct histological areas confirms that our ST-seq data is capable of simultaneously and reliably quantifying spatial gene expression changes in the two species within the same tissue section.

The majority of spots were classified as either human or mouse, however we identified areas of mixed spots where reads mapped to both human and mouse reference genomes. This suggests that these spots contain cells of both human and mouse origin, likely due to the resolution of this platform with each spot comprising multiple cells per spot. Mixed spots were observed in well demarcated tumour-mouse interface regions within each PDOX. Notably, our spatial data clearly define the interface from the human or mouse only regions, which are histologically indistinguishable from surrounding human tumour tissue (Fig. 1b, Fig. S4a). Furthermore, mixed spots were also observed sporadically throughout the bulk of the tissue section irrespective of treatment group, suggesting the infiltration of mouse cells into the bulk of the PDOX, which was independently validated by histological examination. Together, these data validate our spatially resolved transcriptomics workflow and demonstrate our ability to detect discrete tumour and microenvironmental regions within the ST-seq dataset. This highlights the benefits of the ST-approach over traditional sequencing methods that are in capable of delineating intratumoural heterogeneity within a spatial context.

### Spatial transcriptomics identifies and accurately maps cell type subpopulations across the cerebellar cortex and tumour regions of SHH patient-derived MB

To assess data quality and our cell type identification pipeline, we compared unsupervised clustering results of the mouse cerebellar regions with reference mouse brain anatomy. Four distinct clusters were found that overlapped with histologically identifiable regions within the mouse cerebellum (Fig. 2a). To assess if these clusters represented histologically defined cell types, we compared across clusters the expression of well-known gene markers[50] of major cerebellar cell types across clusters (Fig. 2b). Cluster 0 was defined by the high expression of oligodendrocytic or myelin markers (*Mbp,Plp1,Mobp*)[5, 12, 27], consistent with their mapping to expected location within the white matter of the cerebellum. Cluster 1, which mapped to the Purkinje cell layer, had the highest expression of Purkinje cell markers (*Pcp4, Car8, Calb1,Slc1a3, Ppp1r17*)[47]. Cluster 2 was characterised by high expression of granule neuron markers (*Snap25*,*Cbln3*, *Gabra6*)[16, 78], consistent with the observation that this cluster was mapped to the histologically-defined granular layer. Cluster 3 had high expression of *Kit*, a basket/stellate marker that has been demonstrated to identify cells in the molecular layer but not mature Purkinje cells of the cerebellum[1, 16]. This cluster was specifically mapped to the molecular layer in mouse tissue. *Pvalb,* a marker of Purkinje cells whose large dendritic arbour reside within the molecular layer[8, 45], was also highly expressed in this cluster. As an independent validation, without using known markers, we performed gene set activity enrichment analysis for the top ten genes in each of the four clusters (Fig. 2c). Mapping enrichment scores to the tissue, we observed a similarly high concordance of regions with enriched scores with anatomically distinct cerebellum structures. Application of this approach to the other samples yielded similar results (Fig. S2).

To complement marker-based and cluster-based approaches for mapping cell types, we used another independent approach where spots were automatically annotated by using the gene expression profile of the of the dominant cell type and matching this against reference scRNA-seq data from the developing mouse cerebellum [77]. A strong concordance between annotated cell types and known anatomical location of cerebellar subpopulations within the mouse cerebellum was observed (Fig. 2d, Fig. S5). Granule cells and unipolar brush cells were observed in the granule cell layer as previously reported [14, 60], while the molecular layer/ purkinje cell layer were indeed enriched with purkinje cells (Fig. 2d1). As expected, oligodendrocyte precursor cells predominantly mapped to the white matter [75], with astrocytes/ bergmann glial progenitors enriched in the molecular layer/purkinje cell layer consistent with their expected location [11] (Fig. 2d1).

Furthermore, we compared cell types labelled in a blinded fashion from our paediatric neuro-pathologist (TR) to the automated cell types annotated on the basis of dominant cell type as described above using the reference scRNA-seq data from the developing mouse cerebellum [77]. Focusing on the sample with the most extensive annotation, a region labelled as “Mouse macrophage” by our neuro-pathologist was annotated as meninges and microglia (Fig. 2d2), resident macrophages of the central nervous system [42]. Another region labelled as ‘Mouse macrophage and scattered tumour cells’ by our neuro-pathologist was annotated as consisting of astrocytes, microglia, and OPCs (Fig. 2d3) on the basis of ST-seq data. Additional concordance was observed for areas labelled as ‘Meningeal arteries’ by our neuro-pathologist, with ST-seq data further annotating these areas to consist of a number of cell types known to comprise the neurovascular unit surrounding the specialised vasculature of the brain, including meninges, endothelial cells, and pericytes (Fig. 2d4)[42, 55]. Together, these analyses confirm the capacity of ST-seq data and our analytical approaches to define cell types consistent with their expected anatomical location within the mouse cerebellum and with specialist annotation with very high sensitivity and specificity enabling in-depth exploration tumour regions as described below.

### CDK4/6 inhibition reduces tumour heterogeneity

To compare the level of transcriptional heterogeneity within the human tumour sections between treated and untreated samples, we first quantitatively analysed the number of clusters of cells in each sample across a wide range of 23 different clustering resolutions (Fig. 3). The cells that stayed in the same in one cluster or split to two different clusters are shown in the clustertree as arrows (Fig. 3). At the same starting resolution of 0.1, we already observed that spots in the untreated samples split into three clusters, while spots in the two treated samples remained as one or two clusters, suggesting increased heterogeneity in the untreated. We then quantified the extent of the heterogeneity difference by calculating connectivity indexes between clusters [33]. Connectivity is calculated based on the degree that similar spatial spots are in the same cluster. We observed that the two treated samples consistently had much lower connectivity (1171 and 1431) compared to the two untreated samples (2709 and 2029), suggesting less heterogeneity between spatial spots in treated tissues. To quantify tumour heterogeneity between treated and control samples across a wide range of clustering resolutions, we used Shannon entropy and Simpson index [43, 54] which calculate the probability that any two spatial spots belong to one homogenous cluster. Overall, both methods show that treated samples have lower heterogeneity (low Shannon entropy and high Simpson index) compared to control samples (Fig. 3b-d). In addition, comparing the scores at a spot level between the treated and untreated samples showed increased similarity between treated samples (0.02) compared to between the untreated samples (0.04). Overall, the Shannon entropy, Simpson index and connectivity consistently show higher heterogeneity in the untreated samples, indicating that drug treatment reduced the cellular heterogeneity of the tumour.

**Figure 3:**
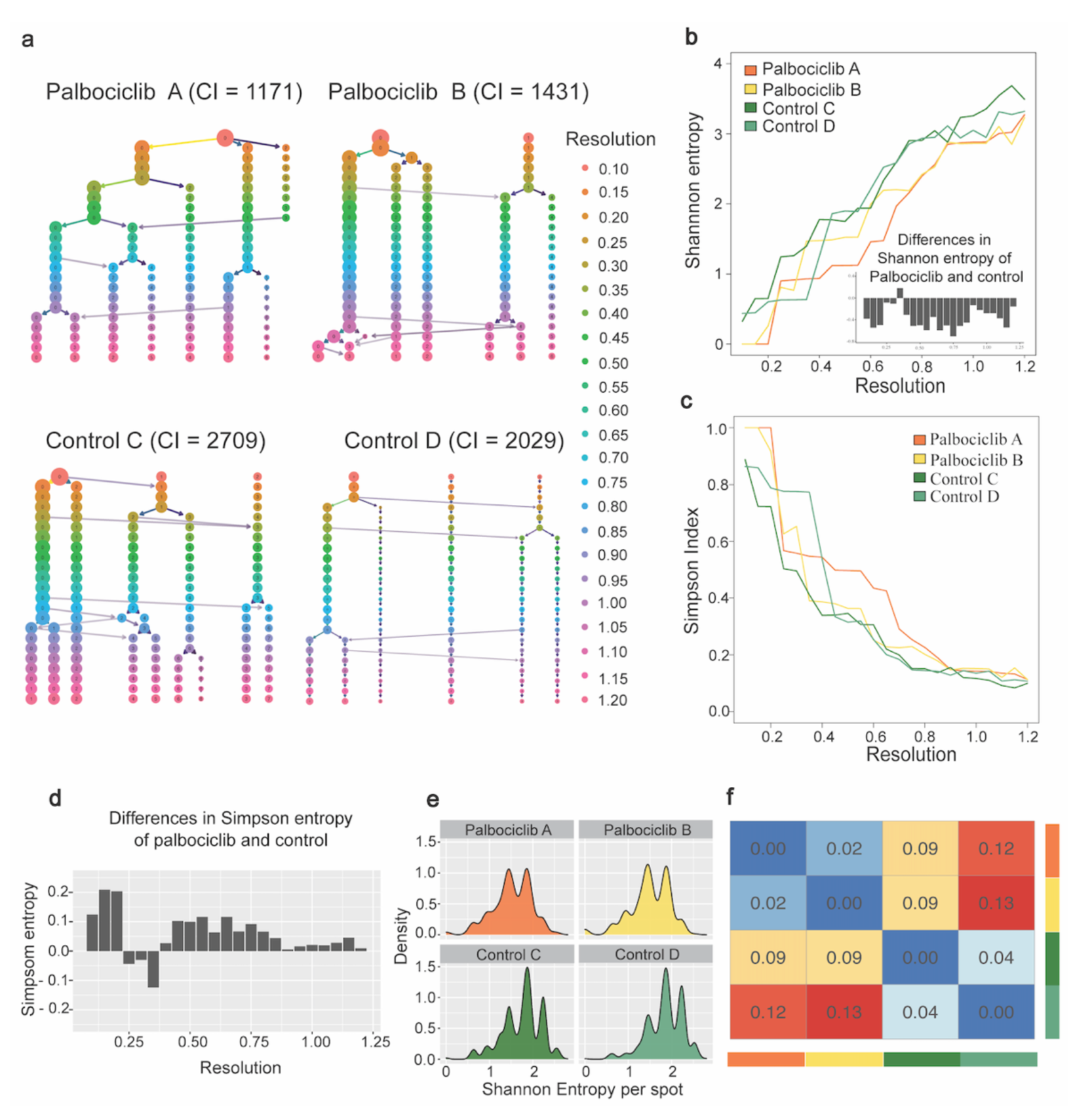
Clustering analysis reveals differences in tumour heterogeneity between Palbociclib treated and control samples. (a) Comprehensive scanning of different clustering resolutions from 0.1 to 1.2 (an interval of 0.05) for each of the four samples. The resolution parameter is used in the graph optimisation procedure to find the optimal number of cell communities (clusters) – a higher resolution leads to more clusters. Cluster partitioning is visualised hierarchically from low resolution (top) to high resolution (bottom), where each row and colour corresponds to the result from one resolution (visualised by Clustree). Branching and merging of cells and clusters are shown by horizontal and vertical arrows when moving from one resolution to the next. CI represents connectivity index, a measure of differences in heterogeneity due to separating or grouping spots that are similar (nearest neighbour). CI values of Controls (2709 and 2029) are higher than Palbociclib (1171 and 1431), indicating the reduced heterogenity in samples under drug treatments. (b) Shannon entropy per sample estimated from each clustering resolution used in a. The x-axis is the same clustering resolution in panel a. The y-axis is the Shannon entropy per sample. Colours represent different individuals. The bar plot on the bottom-right corner shows the differences in Shannon entropy of Palbociclib and control (i.e. mean of entropy in Palbociclib – mean of entropy in control). A high Shannon entropy indicates high heterogeneity, with lower Shannon entropy and heterogeneity in Palbociclib treated PDOX compared to controls *(P<0.001* for Wilcoxson signed-rank exact test and two-tailed paired t-test) (c) Similar to panel b but shows Simpson index. Simpson index represents the probability that two randomly selected spots belong to the same cluster. Thus, a low Simpson index indicates high heterogeneity, with higher Simpson index indicating lower heterogeneity in Palbociclib treated PDOX compared to controls (*P<0.001* for Wilcoxson signed-rank exact test and *P<0.01* for two-tailed paired t-test). (d) Differences in Simpson entropy of Palbociclib and control (i.e. mean of entropy in Palbociclib – mean of entropy in control). (e) Density plot of per spot Shannon entropy using a clustering resolution of 0.8. (f) Pair-wise similarity (i.e. Jensen–Shannon Divergence (JSD)) between distributions in panel E. The same colour bars on the right and bottom indicates the same sample in panel E. The value in the heatmap is the JSD between corresponded samples. The lower the JSD value, the more similar the corresponding distribution.

### CDK4/6 inhibition results in higher levels of neuronal differentiation gene expression in SHH patient-derived MB

To determine the impact of CDK4/6 inhibition on the transcriptome of SHH MB, we compared Palbociclib-treated Med-1712FH and control untreated PDOX across the three spatially defined regions, human, mouse and mixed human mouse interface. The expression levels of >13,000 reliably detected genes were compared. For spots annotated as in human, or mouse or mixed compartments, we applied a pseudobulking strategy to stabilise between-spot variation within each sample, followed by a standard differential expression (DE) analysis using the voom-limma [49] model (Fig. S4b-f). Palbociclib treatment had an effect on all tissue regions, however a more pronounced gene expression change was observed in the human regions compared to the ‘interface’ and mouse regions. A total of 1200 differentially expressed (DE) human genes between Palbociclib-treated and control PDOX were identified in the bulk of the tumour (Fig. 4a, Table S3), compared to 139 DE mouse genes in surrounding mouse tissue (Fig. 4a, Table S5). Moreover, 166 DE human genes were observed when comparing the mixed interface tissue from Palbociclib-treated and untreated PDOX (Fig. 4a, Table S4).

**Figure 4:**
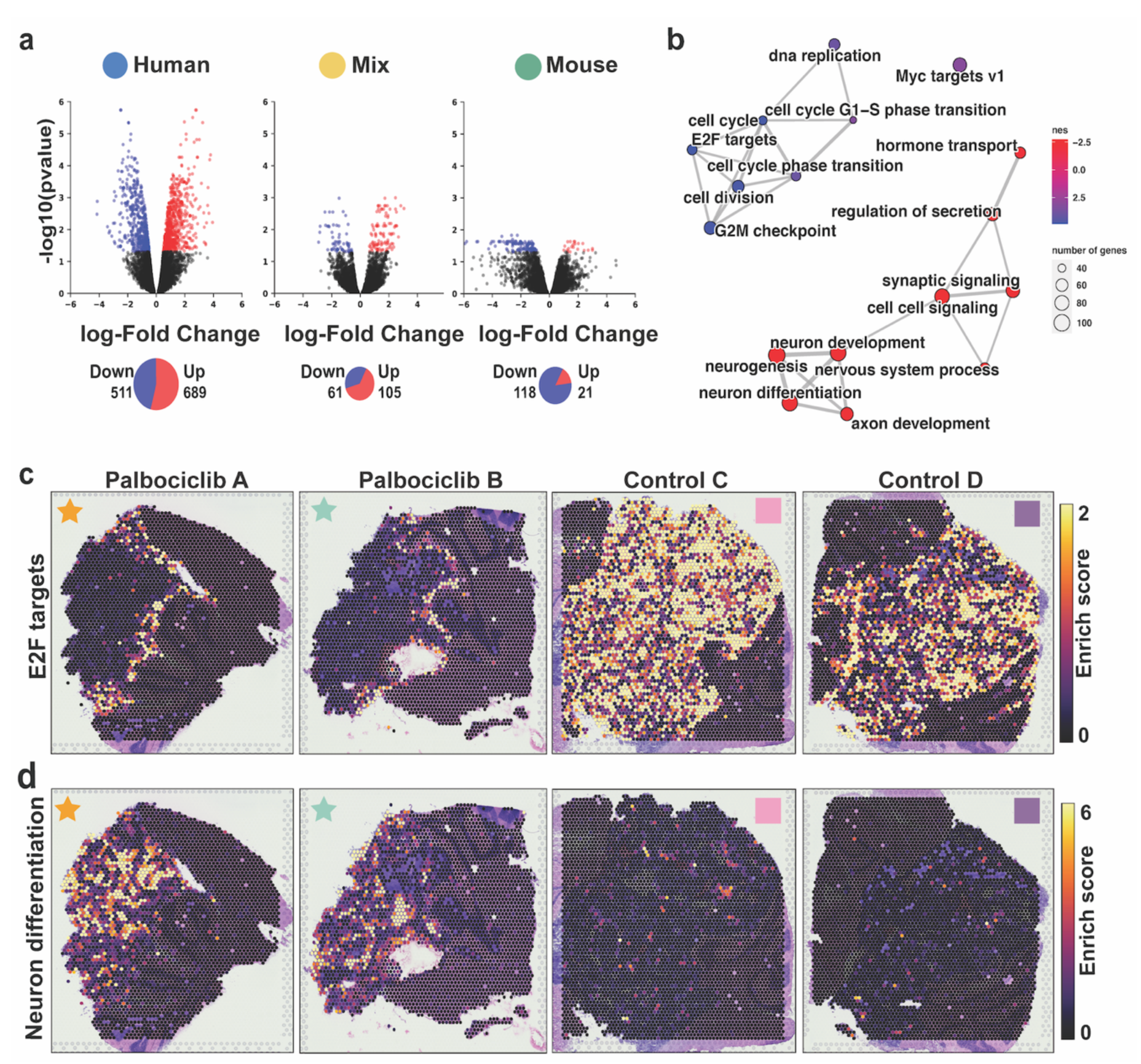
Medulloblastoma downregulate cell cycle and upregulate neuron differentiation related genes in response to Palbociclib treatment in a spatially dependent manner. (a) Volcano plots from analyses of differentially expressed (DE) genes between treatment and control samples, compared for three categories, including human, mix, and mouse spots. DE genes are highlighted with red for upregulated genes, blue for downregulated genes, and black for non-significant change (corrected p < 0.05). Pie charts beneath the volcano plots indicate the number of significant DE genes detected in each comparison. (b) Network visualisation of gene set enrichment analysis results for the human DE genes show upregulation of neuronal differentiation and downregulation of cell cycle. Each dot is annotated with the gene set term. Dot sizes indicate the number of DE genes detected within the term, and dots were coloured according to the normalised enrichment score (NES). Positive NES values (red) indicate upregulation of the term in response to treatment, while negative NES values (blue) indicate downregulation of the term in response to treatment. Edges between terms indicate overlap of the DE genes associated with the connected terms. (c-d) Spot specific gene enrichment scores for *E2F* target genes (c) and neuron differentiation genes (d) are overlaid in the spatial context. High gene set activity is indicated by a bright yellow, and low activity is indicated by dark violet; overall showing continued *E2F* activity at the human-mouse interface in treated samples (c) and neuronal differentiation within the tumour (d). E2F E2 family of transcription factors.

**Figure 5:**
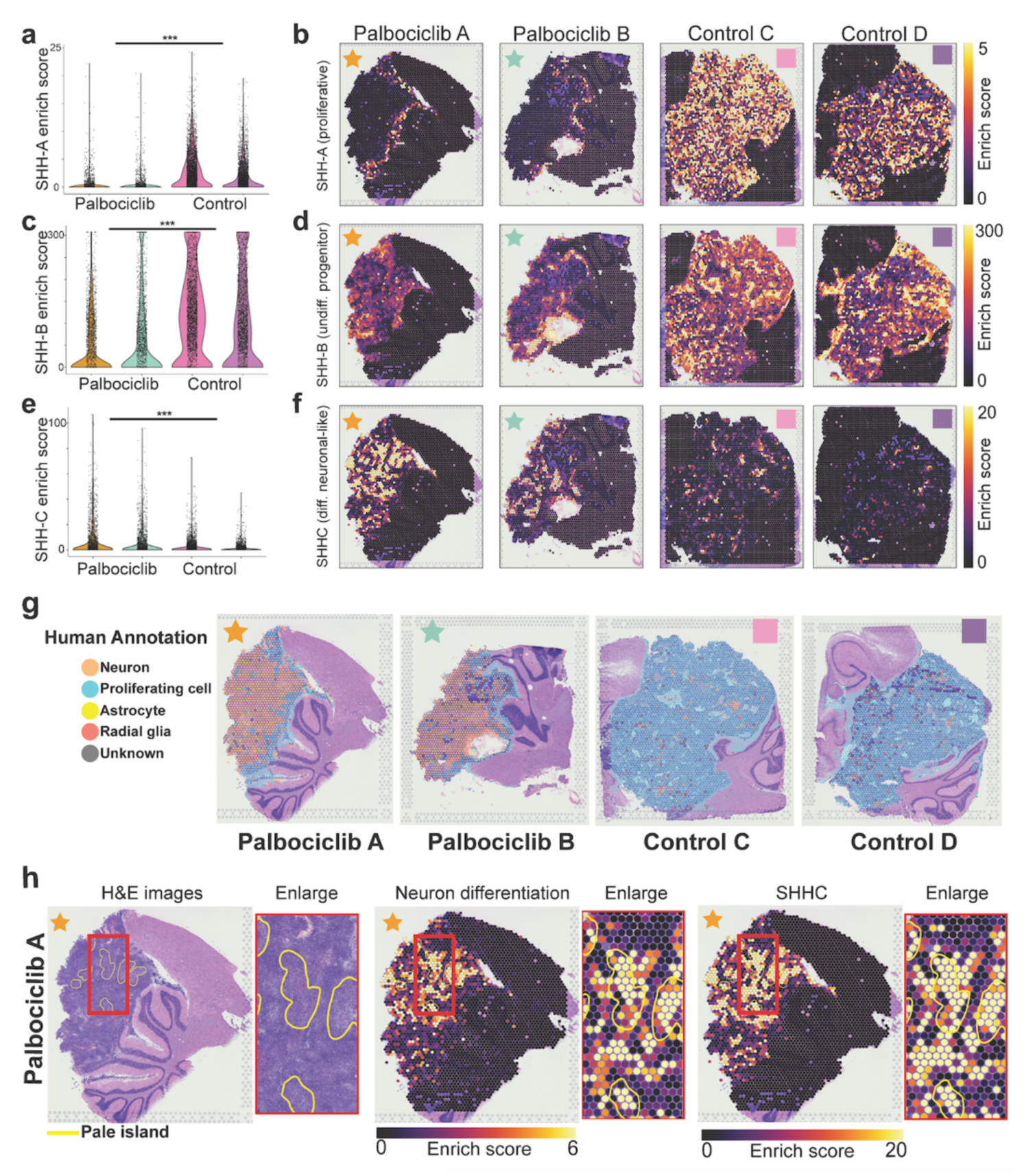
Per-spot gene activity enrichment analysis depicts the spatial localisation of SHH-MB neuronal differentiation, progenitor, and cell cycle in response to Palbociclib. (a) Violin plots of the per-spot SHH-A (cell cycle) gene signature activity, separated by Palbociclib A, Palbociclib B, Control C and Control D samples (left to right). *** indicates p-value <0.001 when comparing these scores between Palbociclib-treated and control samples using a pseudosampling approach with two tailed t-test (see Methods). (b) SHH-A (cell cycle) gene signature enrichment scores in Palbociclib A, Palbociclib B, Control C, and Control D samples in their spatial context. Spots are coloured according to enrichment scores, from bright yellow (high gene set activity) to purple (low gene set activity). (c-f) Each row of panels is equivalent to the first row described above (panels a-b), except the middle row (panels c-d) indicate enrichment scores for the SHH-B progenitor cell signature, while the bottom row (panels e-f) indicates enrichment scores for the SHH-C neuronal differentiated signature. Both of these latter signatures also showed significant difference (indicated by ***), with SHH-B showing significant downregulation and SHH-C showing significant upregulation in response to Palbociclib treatment. (g) Automated annotation of human/mix spots using pattern matching against a reference scRNA-seq dataset from 13 week old male foetal human brain. Annotations indicate proliferative cells at the tumour human-mouse interface in treated samples with neuronal cells at the tumour core, while proliferative cells are observed throughout untreated samples. (h) Comparision between pathologist annotations from H&E images (left) and per-spot enrichment of DE neuron differentiation genes and SHH-C (neuronal) signature subpopulation in Palbociclib A. Yellow circles indicate the ‘pale island’ regions annotated independently by a pathologist based on histology.

Palbociclib down-regulated genes were statistically enriched for Molecular Signature Database (mSigDb) Hallmark and Biological Process Gene Ontology (GO) terms related to cell cycle pathway (e.g. E2F targets (genes *ATAD2, MCM2, BIRC5*), regulation of cell cycle, cell cycle G1/S phase transition), mitosis (e.g. mitotic spindle organisation, regulation of mitotic nuclear division, regulation of mitotic cell cycle and mitotic nuclear division) and microtubule organisation (e.g. microtubule organizing centre organisation, microtubule cytoskeleton organisation) (Fig. 4b, Table S6). GO and mSigDb enriched terms for up-regulated genes following drug treatment include neurogenesis (e.g. neuron differentiation (genes *STMN2, STMN4, TUBB3*), regulation of neurogenesis and neuron development), cell-cell signalling (e.g. synaptic signalling) and ion transport (e.g. cation transmembrane transport). We sought to validate the higher levels of neuron differentiation observed in response to Palbociclib treatment by immunostaining using STMN4 and TUBB3. A significant increase in both STMN4 (*P<0.05*) and TUBB3 (*P<0.05*) expression was observed throughout the majority of treated SHH PDOX (Fig. S8), confirming the up-regulation of neuron differentiation identified with our ST-seq data. Further in-depth and independent gene set activity enrichment analysis and cell type assignment at spot level across the tissue, as described later in further detail below, support the differential regulation of cell cycle and neuronal differentiation by Palbociclib.

Down-regulation of *E2F* target genes in drug-treated PDOX is expected given Palbociclib blocks *E2F* transcription factor activity and subsequent cell cycle progression[29]. On this basis, we hypothesised that *E2F* targets could be used as a spatial surrogate of Palbociclib activity across the intact SHH PDOX. To spatially define the effect of Palbociclib, we performed per-spot gene enrichment analysis of the activities of *E2F* target genes and compared these activities in drug-treated and control PDOX. As expected, we observed a contrasting pattern of *E2F* pathway activity between untreated and treated PDOX. High *E2F* pathway activity was observed throughout the entire tumour of untreated SHH PDOX, with this high *E2F* activity restricted to the ‘interface’ region only in drug-treated PDOX (Fig. 4c). Using the same approach, we investigated whether the enrichment of neuronal differentiation occurred in a spatial context. Whilst control PDOX contained small islands of tumour cells with high levels neuronal differentiation gene activity, these islands were much larger and comprised the majority of the human tumour following Palbociclib treatment (Fig. 4d). Taken together, these data are consistent with our DE analysis (Fig. 4a) and suggests a spatial response of SHH PDOX to Palbociclib treatment, with the drug appearing to have a limited effect on the interface region.

### Tumour cells at the tumour-microenvironment interface continue to proliferate despite Palbociclib treatment

Our DE analysis suggests distinct biological processes are operating at the tumour-microenvironment interface region, where tumour cells continue to proliferate despite Palbociclib treatment. To further characterise our observed spatial response to Palbociclib treatment, we used transcriptional signatures generated by scRNA-seq of SHH MB patient samples[36]. A subset of 100 genes from three transcriptional signatures, cell cycle activity/proliferation (SHH-A), undifferentiated progenitors (SHH-B), and neuronal-like programs (SHH-C)[36] were used for this analysis (Fig. 5a-f). Spatial enrichment of the cell cycle SHH-A transcriptional state strongly overlapped with the spatial pattern observed for E2F-target gene activity. High activity of SHH-A was observed throughout the tumour region of untreated SHH PDOX, with this expression pattern once again restricted to the interface region of treated PDOX (Fig. 5a). This is of particular relevance considering the limited overlap between these two gene lists, with only 34/100 in SHH-A shared with the E2F pathway. We went on to confirm this spatially-restricted response in drug-treated PDOX using immunofluorescence analysis for cell proliferation marker, Ki67. The proportion of proliferative cells quantified in the control compared to the bulk and interface regions of drug-treated PDOX validated that observed at the transcriptional level (Fig. S9), with 58.95% of cells Ki67+ in the untreated SHH PDOX compared to 6.86% in the bulk region of drug-treated PDOX (*P<0.0001).* The proportion of proliferative cells in the interface of drug-treated PDOX however remained at very similar levels to untreated PDOX, with 43.37% of cells Ki67+, confirming the continued proliferation of tumour cells at this region.

The undifferentiated progenitor SHH-B signature was observed at lower levels in drug-treated PDOX (Fig. 5c), with high expression of SHH-B still observed in foci throughout the tumour, suggesting the retention of small pockets of undifferentiated progenitors in drug-treated tumours (Fig. 5d). In contrast, the neuronal-like SHH-C transcriptional program was more highly expressed in Palbociclib-treated tumours compared to untreated PDOX (Fig. 5e,f). High expression of the SHH-C signature was observed in large ‘islands’ throughout treated samples, consistent with the spatial enrichment pattern of the neuronal differentiation pathway. Only 15/100 SHH-C genes overlapped with the neuronal differentiation pathway, again suggesting almost completely independent validation. Calculating gene set activities for SHH-B and SHH-C signatures per-spot across all tumours, significantly higher density values of the SHH-C signature (*P<0.001*) and lower density values of the SHH-B signature (*P<0.001*) were identified following Palbociclib treatment (Fig. S10). We also found that areas with high per-spot enrichment of SHH-C and neuronal differentiation correlated with areas annotated by our neuropathologist as “pale islands” (Fig.5h, Fig. S11). This indicates the correlation of neuron differentiation with nodule formation and desmoplasia previously associated with a favorable prognosis in MB [57, 70]. Together, our spatial gene set activity analysis confirms findings from DE analysis, whereby Palbociclib treatment increases the expression of genes associated with neuronal differentiation and reduces cell proliferation in the bulk of the PDOX, with very little drug impact at the interface region.

To further validate these findings, we performed automated cell type annotation of spots assigning dominant cell types to each spatial tissue spot, based on gene expression correlation between our ST-seq data with an additional public reference scRNA-seq data from human foetal brain samples [2, 32] (with a focus on proliferative cell types). Conversely, treatment of SHH PDOX resulted in a shift in this proliferation and differentiation (neuronal) states, with drug-treated PDOX comprising almost entirely of neurons within the bulk of the tumour with the proliferating cell type 1 annotated only at the mixed tumour interface (Fig. 5g). Together, spatial gene expression analyses conclusively demonstrate that the MB/microenvironment interface continues to proliferate despite Palbociclib treatment, indicating the importance of the TME in regulating response to therapy. Our analysis strategy incorporating ST-seq technologies with information from scRNA-seq on SHH signatures and brain cell types clearly elucidates the complexity and heterogeneity of the MB TME and highlights how these interactions strongly influence response to therapy.

### Astrocytes are the most dominant cell type present at the tumour-microenvironment interface

Our ST-seq results define a transcriptionally distinct interface region where tumours contact the microenvironment and continue to proliferate despite Palbociclib treatment. We next sought to identify what cell types reside within the microenvironment that may be influencing the response of tumour cells to Palbociclib. To do this, we revisited our correlation between manual pathologist annotation and ST-seq spot annotation on the basis of dominant cell type using correlation-based approach to map (transfer) labels from a reference scRNA-seq dataset [77] to each spatial spot (Fig. 2d, Fig. S5,6).

Having previously established accurate spot cell type annotations on the basis of ST-seq data (Fig. 2d, Fig. S5,6), we next sought to determine what mouse cell types may be residing at this interface region. Using a mouse reference dataset of developing mouse cerebellum [77], we annotated cell types for mouse spots at the border. Astrocytes and Bergmann glia were identified as the most abundant mouse cell type observed at the border of the interface region, clearly abutting tumour cells in both untreated and drug-treated PDOX (Fig. 2d, Fig. S5,6). As our approach to predict glial cells was consistent with independent pathological annotation, we further explored enrichment of glial cells across the tissue using both cell type annotation and marker-based expression visualisation. We observed the enrichment of Astrocyte/Bergmann glia in the tumour invasion region, largely overlapping the interface region (Fig.6a-b, Fig. S12,

Table S7, p value<0.01). We next applied statistical methods to compare the unsupervised annotation results with Astrocyte/ Bergmann glia marker, *Gfap* (Fig. 6b). We observed a consistent trend of *Gfap* expression associated with both the tumour border region and the spots previously annotated as Astrocyte/Bergmann glia. Finally, we sought to validate the presence of proliferating tumour cells with Astrocyte/Bergmann glia in Palbociclib-treated PDOX by immunofluorescence using Gfap as well as Ki67 to mark proliferating cells. Immunofluorescence of both untreated and drug-treated PDOX showed strong Gfap expression throughout the surrounding mouse brain tissue (Fig. 6c), confirming the presence of Astrocyte/Bergmann glia throughout this tissue. Untreated tumours were comprised of almost entirely of proliferating cells, however a clear reduction in proliferating cells was observed in drug-treated tumours, with proliferating cells highly associated with the Gfap-positive surrounding mouse brain (Fig. 6c). We sought to investigate whether astrocytes associated with proliferative tumour cells within the tumour-microenvironment interface were distinct to those within the mouse only regions of the brain. Using ST-seq data, DE analysis identified that the astrocytes in the interface and in the mouse only region are molecularly distinct and respond differently to Palbociclib treatment (Fig.S13). Through unbiased cell-cell interaction analysis, we found enriched interactions of astrocyte-microglia at the interface (Figure S14). Together, using multiple independent approaches, we confirmed the enrichment of mouse astrocytes and glia localised to the interface region, suggesting their possible role in the differential response of this region to Palbociclib treatment.

**Figure 6:**
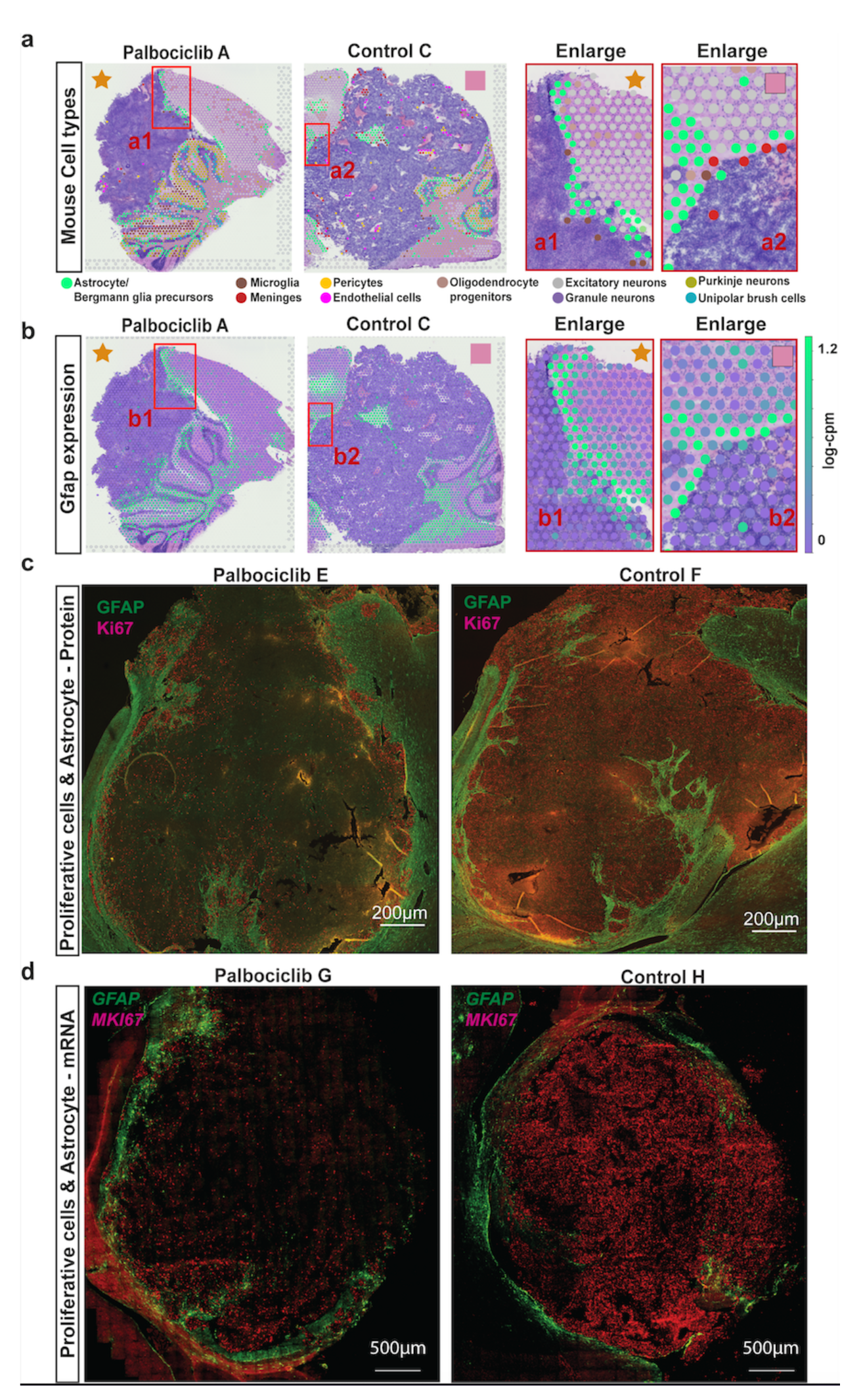
Astrocytes are localised to the mixed tumour-microenvironment interface. (a) Palbociclib A and Control C shown with annotated mouse cell types depicting the localisation of astrocytes to the interface region. Spot colours indicate cell types. Red boxes indicate the region of interest enlarged on right. (b) The equivalent to (a) but with expression of the astrocyte marker genes *Gfap*. The gradient colour reflects log cpm (count per million). A strong concordance between astrocyte localisation and Gfap expression is observed, with both localised at the interface region. (b) The reactive astrocytes (Gfap) located in the tumour–mouse interface. (c) Positive proliferating cells (Ki67) are located throughout the tumour in untreated tumours, but localised to the tumour–mouse interface in drug-treated PDOX in close proximity with astrocytes, as identified by GFAP staining. (d) Target RNA molecule expression of astrocytes (*Gfap*) and proliferative tumour cells (*MKI67*) at a single cell level using RNAscope.

To further investigate the possible role of mouse astrocytes and glia in coordinating the differential response to Palbociclib, we next performed spatial transcriptomic analysis via smRNA-FISH/RNAscope on three representative samples of untreated and drug-treated PDOX. We multiplexed eight genes to validate astrocytes and microglia as identified on the basis of ST-seq and immunofluorescence staining and their specific localisation with respect to proliferating tumour cells within the tumour microenvironment interface at single cell resolution. Using *MKI67* as a marker of proliferating tumour cells, we validated the spatially-restricted response of drug-treated PDOX to Palbociclib (Fig. 6d). High *MKI67* signal was restricted to the tumour cells of the interface located in close proximity to *Gfap*-positive astrocytes, which were completely absent in the bulk of drug-treated PDOX comprising *MKI67*-negative tumour cells (Fig. S15). This localisation pattern is consistent with ST-seq and immunofluorescence staining results suggesting again a role for astrocytes in the continued proliferation of tumour cells within the interface. Recently, one study reported that Il-4 secreted from astrocytes stimulates Igf1 production from brain-resident microglia to promote progression of SHH-activated mouse MB[82] (Fig. 7a). We confirmed *Cd68/Aif1-*expressing tumour-associated microglia and *Gfap/S100b*-expressing astrocytes in close proximity to proliferating (*MKI67*-positive) tumour cells within the interface of drug-treated PDOX (Fig. 7b-c). While *Il4r* and *Igf1* mRNA was expressed in *Cd68/Aif1-*positive tumour-associated microglia throughout drug-treated PDOX (Fig. 7c-d), *Il4* expression co-localised with *Gfap/S100b*-expressing astrocytes exclusive to the interface region of drug-treated PDOX. Taken together, these data revealed that the tumour microenvironment differs between areas of differential drug response, with correlative evidence for a multicellular paracrine feedback loop involving astrocytes and tumour-associated microglia promoting the continued proliferation of tumour cells in the interface of drug-treated PDOX.

**Figure 7.**
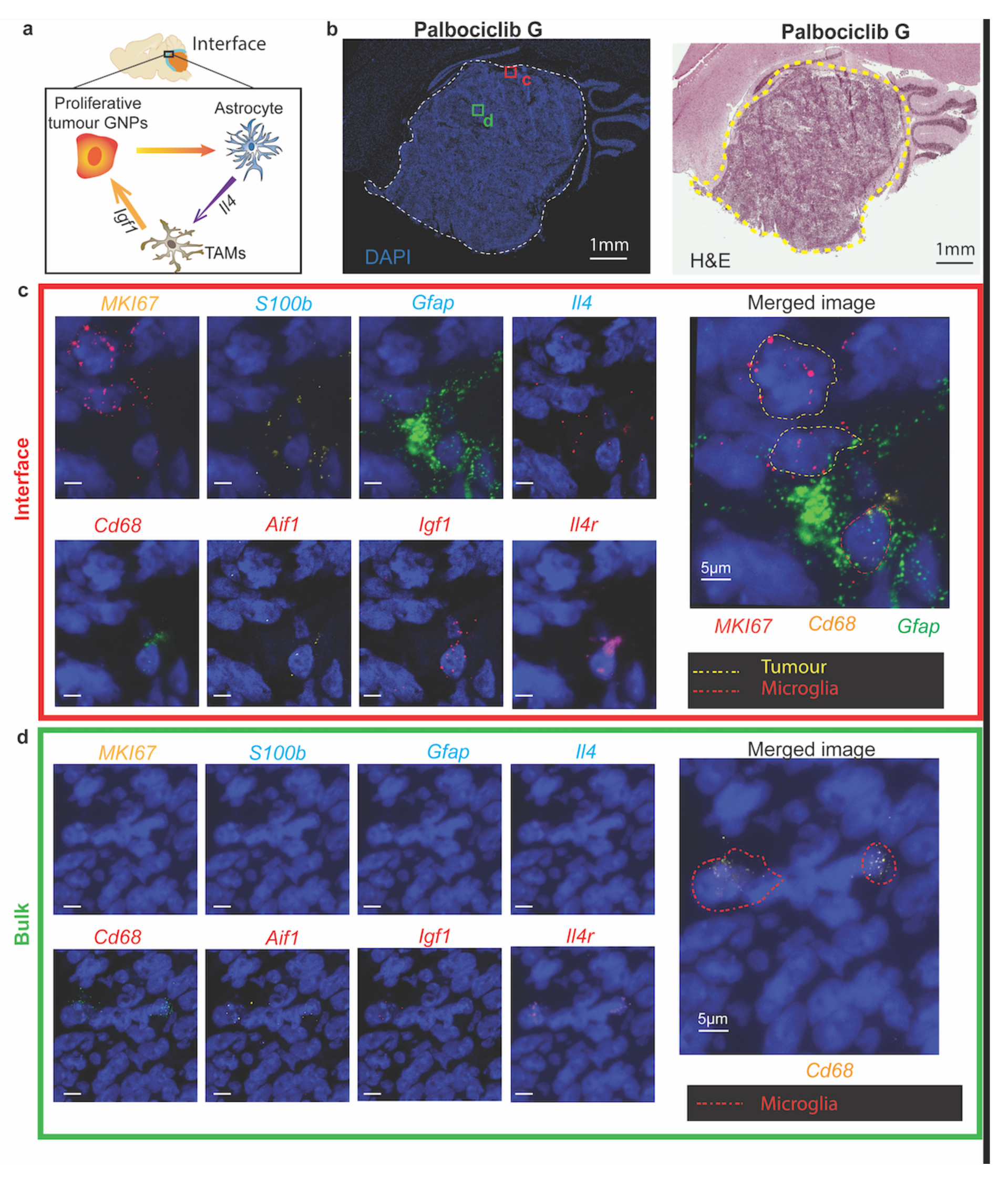
The identification of a multi-lateral network associated with continued tumour cell proliferation in within the drug-treated tumour microenvironment interface. (a) Proposed hypothesis facilitating continued proliferation of tumour cells at the interface, a consequence of interactions between proliferative tumour GNPs and host cell types such as microglia and astrocytes. (b) DAPI and H&E image overview of a drug-treated tumour, Palbociclib G. (c) Representative target RNA molecule expression at a single cell level using RNAscope for gene markers of cell types and associated ligands and receptors of proposed hypothesis: microglia (*Cd68, Aif1, Igf1, Il4r*), astrocytes (*S100b, Gfap, Il4*) and proliferating tumour cells (*MKI67*) within the tumour microenvironment interface and (d) tumour bulk. GNPs Granule Neuron Progenitors, TMAs Tumour Associated Microglia. Scale bar: 5um.

## Discussion

New targeted therapies are urgently needed for children diagnosed with MB. However, effective treatment is met with several challenges due to, but not limited to, the extensive heterogeneity of the disease. Significant genomics, epigenomics and transcriptomics efforts have facilitated a comprehensive understanding of the molecular basis underpinning MB [35, 41, 51, 61], with the well-appreciated intertumoural heterogeneity recognised by the consensus of up to 12 biologically and clinically relevant subtypes of disease. Recent single cell RNA sequencing approaches have begun to characterise the intratumoural heterogeneity of MB, defining diverse neoplastic [36, 77] and subgroup-specific stromal and immune cell subpopulations [70] comprising individual tumours. These studies have advanced our understanding of the inherent intratumoural cellular heterogeneity of MB and are beginning to address the contribution of the broader TME in MB, however the spatial organisation and interactions of these cells within tumours still remains poorly understood. Spatial microdissection of MB biopsies followed by standard RNA sequencing previously identified different cancer clones within a tumour, but failed to identify cell-types within a clone or map cellular distribution within their microenvironment [59]. Here, we used spatially resolved transcriptomics to define the transcriptional state and location of tumour cells within intact tissue sections of a PDOX model of SHH MB. We generate spatial molecular maps for this PDOX model of SHH MB and integrate this with scRNA-seq data from human MB patients to assess how this intratumoural heterogeneity changes following treatment with the CDK4/6 inhibitor Palbociclib. Our study provides several fundamental insights and highlights the importance of considering both the molecular and spatial context of all cell types when assessing the efficacy of any therapeutic approach in neuro-oncology.

Intratumoural heterogeneity is a key determinant of therapeutic resistance (Reviewed in Jamal-Hanjani, Quezada, Larkin and Swanton [38]), with a better understanding of cellular drug response patterns critical for future development of more effective combination therapies. Our analysis reveals that SHH PDOX contain multiple malignant transcriptional states, recapitulating the extent of intratumoural cellular diversity identified in primary human SHH MB [36, 77] . This finding, together with others [7, 85], confirms the translational relevance of PDOX as preclinical model systems and supports their use for interrogating the molecular and cellular basis of intratumoural heterogeneity underlying therapy response. Here we have defined that Palbociclib treatment depletes the diversity of transcriptional states within the SHH

PDOX model of MB, consistent with a previous scRNA study treating a SHH GEMM models of MB with the SHH inhibitor, Vismodegib. This reduction in cellular diversity that we and others observe following therapy may be advantageous for the treatment of residual tumour cells following therapy, as long as biological pathways or processes are shared between these remaining cell types and can be therapeutically targeted.

Our study offers new insight into the previously described features of CDK4/6 inhibition in MB, with Palbociclib treatment inducing neuronal differentiation across all remaining cell types comprising the bulk of the SHH PDOX. This was also correlated to elevated neuronally differentiated SHH-C transcriptional state from human MB scRNA studies. Understanding the mechanisms that control this state of neuronal differentiation in response to CDK4/6 inhibition could facilitate the identification of therapies to eradicate this residual disease. Cyclin D1 was recently shown to cooperate with CDK4 to directly phosphorylate and upregulate Atonal homolog 1 (Atoh1), a transcription factor indispensable for maintaining granule neuron progenitors, the cells of origin of SHH MB [77], in their immature state [58] . This suggests that Palbociclib treatment may not only be halting the cell cycle, but also blocking the CDK4/CyclinD1 kinase-mediated phosphorylation of *ATOH1* leading to neuronal differentiation. One additional study outlines a role for chromatin remodelling following Palbociclib treatment, increasing levels of several activator protein (AP)-1 transcription factors which drive enhancer activity and upregulate luminal differentiation gene signatures resulting in mammary tubule formation indicative of a more differentiated state breast cancer tissue [79]. Other enhancers were found to govern apoptotic evasion, lending support for therapeutic combinations comprising CDK4/6 inhibitors and Bcl-xL inhibitors. Future studies are required to dissect the possible regulation of ATOH1 and epigenome changes as a consequence of CDK4/6 inhibition in SHH MB and whether this differentiated state is terminal or cells retain the capacity to de-differentiate. Such data are urgently required for the identification of potential therapeutic combinations that may eliminate this residual differentiated neuronal population.

Spatial mapping technologies have helped build upon single cell studies demonstrating that cell populations residing in distinct TME regions display differential therapeutic responses [3]. Our analysis is consistent with this, identifying a spatially distinct transcriptional state at the tumour-microenvironment interface, similar to that previously described in melanoma [37], which did not respond to Palbociclib treatment. We confirmed this finding by mapping transcriptional states derived from scRNA-seq studies of human SHH to our PDOX SHH model, with the proliferative SHH-A transcriptional state mirroring elevated *E2F* pathway activity in this region. Our previous work investigating blood-brain-tumour-barrier integrity of this tumour model suggests that the lack of response in this tumour region may be in part a consequence of insufficient drug exposure, with an intact blood-brain-tumour-barrier (BBTB) identified in tumour tissue abutting the surrounding brain parenchyma [24]. This is supported by an additional study which demonstrated that therapeutic sensitivity of MB was previously correlated to presence of a disrupted BTBB [66] . Future studies aimed at quantitating the spatial tumour tissue drug concentrations and correlating this to pharmacodynamic readouts of drug efficacy using ST-seq are required to determine whether inadequate drug delivery is responsible for the lack of drug effect in this interface region.

Continued proliferation in this interface region may also be due to crosstalk between tumour cells and TME cells of the surrounding parenchyma. Indeed, spatially confined microenvironmental states within human pancreatic cancer have been shown to execute distinct tumour-promoting and chemoprotective functions [28], also very well recognised in Glioblastoma [4, 9, 26, 76]. We clearly show that both mouse and human cells precisely map to the interface region that continues to proliferate despite Palbociclib treatment, suggesting that the lack of drug response in this interface region could also be due to the interactions between MB cells and non-tumour cells of the surrounding parenchyma. Through the analysis of mouse cell types residing in that region, we show that astrocytes and microglia were the predominant cell types residing in this interface region. Astrocytes are increasingly recognised as playing an indispensable role in the progression of SHH MB [13, 30, 52, 83], with one recent study further defining a multi-lateral network within the TME where crosstalk between tumour-associated astrocytes and microglia promotes SHH tumour progression [83]. The significant abundance of both these cell types in this interface region suggests that the continued tumour proliferation here despite Palbociclib treatment may also be due to crosstalk with these two cell types. These findings support further studies focused on interactions between cell types of this multi-lateral network, to investigate whether this may be relevant to other treatment modalities and to identify potential therapeutic targets that may be exploited.

## Conclusion

Our study is, to the best of our knowledge, the first spatially resolved gene expression atlas of SHH PDOX MB and acts as proof-of-principle for the use of ST-seq in identifying spatially organised tumour heterogeneity of MB. These data provide further insight into the intratumoural heterogeneity of SHH MB and highlight the importance of considering both the molecular and spatial context of cell types when assessing the efficacy of any therapy. Our findings have important implications for the mechanisms of efficacy and resistance for CDK4/6 inhibitors in MB, but also more broadly for the preclinical and clinical assessment of any therapeutic approach in neuro-oncology. The advent of more recent ST-seq platforms with increased single cell resolution combined with multiOMIC approaches and temporal analysis of several PDOX MB models will provide a greater understanding of the tumour response to therapy.

## List of abbreviations

### MB – Medulloblastoma; SHH – Sonic Hedgehog; CDK4 – Cyclin Dependent Kinase 4; CDK6

-Cyclin Dependent Kinase 6; WNT – Wingless; Gp3 – Group 3; Gp4 - Group 4; IHC – immunohistochemical; scRNA-seq - single cell RNA sequencing; TME - tumour microenvironment; ST-seq - Spatial transcriptomics sequencing; E2F – E2 family of transcription factors; PDOX - patient-derived orthotopic xenograft; *Mbp –* Myelin Basic Protein*; Plp1 –* Proteolipid Protein 1*; Mobp –* Myelin Associated Oligodendrocyte Basic Protein*; Pcp4 –* Purkinje Cell Protein 4; *Car8 –* Carbonic Anhydrase 8; *Calb1* – Calbindin 1; *Slc1a3 -* Solute Carrier Family 1 (Neutral Amino Acid Transporter) Member 3; *Ppp1r17 –* Protein Phosphatase 1 Regulatory Subunit 17; *Snap25* – Synaptosome Associated Protein 25; *Cbln3* – Cerebellin-3; *Gabra6 –* Gamma-Aminobutyric Acid Type A Receptor Subunit Alpha6; *Kit –* KIT Proto-Oncogene, Receptor Tyrosine Kinase; *Pvalb –* Parvalbumin; *ATAD2 –* ATPase Family AAA Domain Containing 2; *MCM2 –* Minichromosome Maintenance Complex Component 2; *BIRC5 –* Baculoviral IAP Repeat Containing 5*; STMN2 –* Stathmin 2; *STMN4 -* Stathmin 4; *TUBB3 –* Tubulin Beta 3 Class III; DE – Differentially expressed; *Gfap –* Glial Fibrillary Acidic Protein; *ATOH1 –* Atonal BHLH Transcriptional Factor 1; BTBB - blood-brain-tumour-barrier.

## Declarations

### Ethics approvals and consent to participate

All experimental protocols were approved by The University of Queensland Molecular Biosciences committee. This study involved no human participants. The procedures used in this study adhere to the tenants of the Declaration of Helsinki.

## Consent for publication

Not applicable as study using de-identified data.

## Author contributions

L.A.G designed the research project, L.A.G and Q.N designed research methods, T.V, K.J, J.C, A.M, E.T, T.R and L.A.G performed research, B.B, T.V, G.N, D.D.B, M.J.D, Q.N, contributed new datasets, reagents and analytical tools, T.V, B.B, Q.N, L.A.G. analysed the data, T.V, B.B, K.J, Q.N, B.J.W and L.A.G. wrote the manuscript. All authors have reviewed and approved the final version of the manuscript.

## Acknowledgements

The authors would like to thank the patients and families who contributed brain tumour tissue for the PDOX model. This research was carried out in part at the Translational Research Institute, Woolloongabba, QLD, 4102, Australia. Samples were sequenced by Institute for Molecular Bioscience sequencing facility. We thank the participants of this facility and acknowledge the contributions of the investigators to this study.

## Supplementary Tables

**Supplementary Table S1**: Clustering analysis to identify gene markers distinguishing cerebellar structures within the mouse brain compared to known mouse brain anatomy

**Supplementary Table S2**: Species cutoffs to stratify spots as human, mouse or mix in each sample

**Supplementary Table S3**: Differentially expressed genes within the human spatial region of the tissue as a consequence of Palbociclib treatment.

**Supplementary Table S4**: Differentially expressed genes within the mix spatial region of the tissue as a consequence of Palbociclib treatment.

**Supplementary Table S5**: Differentially expressed genes within the mouse spatial region of the tissue as a consequence of Palbociclib treatment.

**Supplementary Table S6**: Gene Set Enrichment Analysis of the human differentially expressed genes in Palbociclib treated tumours compared to untreated control tumours.

**Supplementary Table S7**: Descriptive statistics supporting cell type enrichment at the tumour-brain interface for each sample.

## Supplementary Figures

**Supplementary Figure S1.**
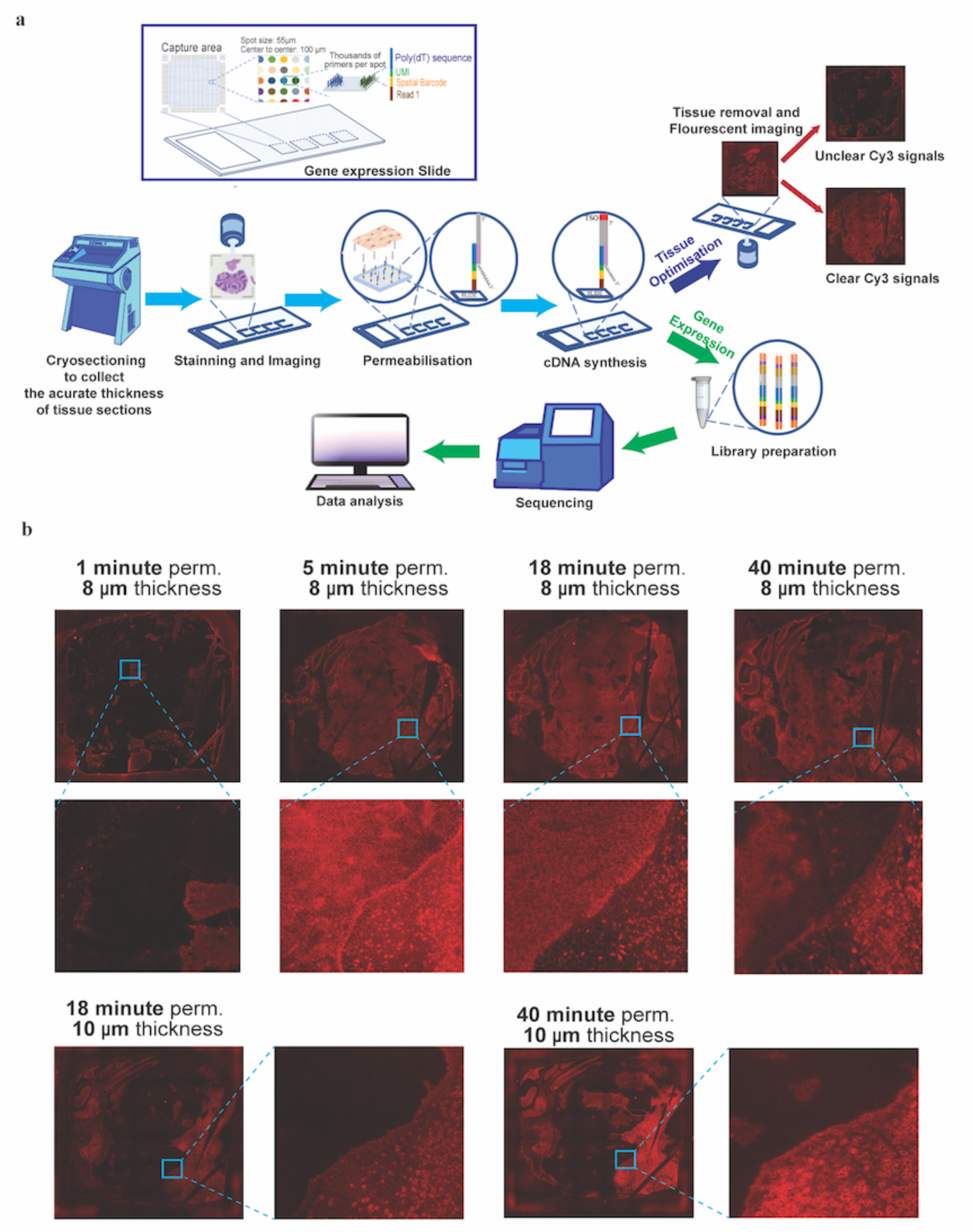
Optimisation of the spatial Transcriptomics (Visium) technology for hybrid human-mouse tissue sections. A. Visium array. Top panel shows the design of an array-printed glass slide (Visium, 10x Genomics). Four capture areas (size of 6.5mm x 6.5 mm) are shown, each contains -5,000 unique spatially barcoded spots. Spot size is 55μm with 100 μm centre to centre distance. Each spot has millions of probes with poly(dT) sequences, UMI, spatial barcode sequence, and lllumina Truseq read 1 adapter. B. Visium workflow. Bottom panel shows visium workflow. Prior to library preparation (gene expression assay - GEx assay), Tissue Optimisation (TO assay) was performed to optimise permeabilisation conditions that worked for both human and mouse cells. In the TO step, tissue samples are sectioned, placed onto the microarray slide, fixed, stained, permeabilised. The released RNA is used for cDNA synthesis with fluorescence labelled nucleotides and the slide is imaged by microscopy to check for the mRNA capture efficiency. Based on the quality of the fluorescence signal across the whole sections, we found the optimal sectioning thickness and permeabilization time for the mouse PDOX system (described in B). In the actual gene expression GEx process, the library preparation is prepared after cDNA synthesis, followed by sequencing and data analysis. The cDNA is synthesised from the captured RNA molecules directly on the slide, and then released from the microarray and collected. The cDNA later is used to generate a sequencing library. Sequencing reads contain spatial information based on the unique spatial barcodes. Consequently, the gene expression measurements can be visualised and analysed together with their originated locations within the tissue and can be mapped to the high-resolution tissue image. C. Tissue optimisation of MB-PDX. Distribution of Cy3-labelled cDNA signals (strong red suggests high signal; dark means low/no signal) across the human and mouse tissue sections were. For tissues sectioned at 10μm, across a range of permeabilisation time-points from 1 minute to 40 minutes, we consistently found that large areas, especially the tumour in the centre, showed no cDNA signal (dark regions). For tissues sectioned at 8μm we tested permeabilisation time-points ranged from 1 minute to 40 minutes. Except for the 1-minute permeabilisation option, other time-points showed cDNA signals across the entire tissue; however, the signal at 40 minute permeabilisation was highly diffused, suggesting over-permeabilization.

**Supplementary Figure S2.**
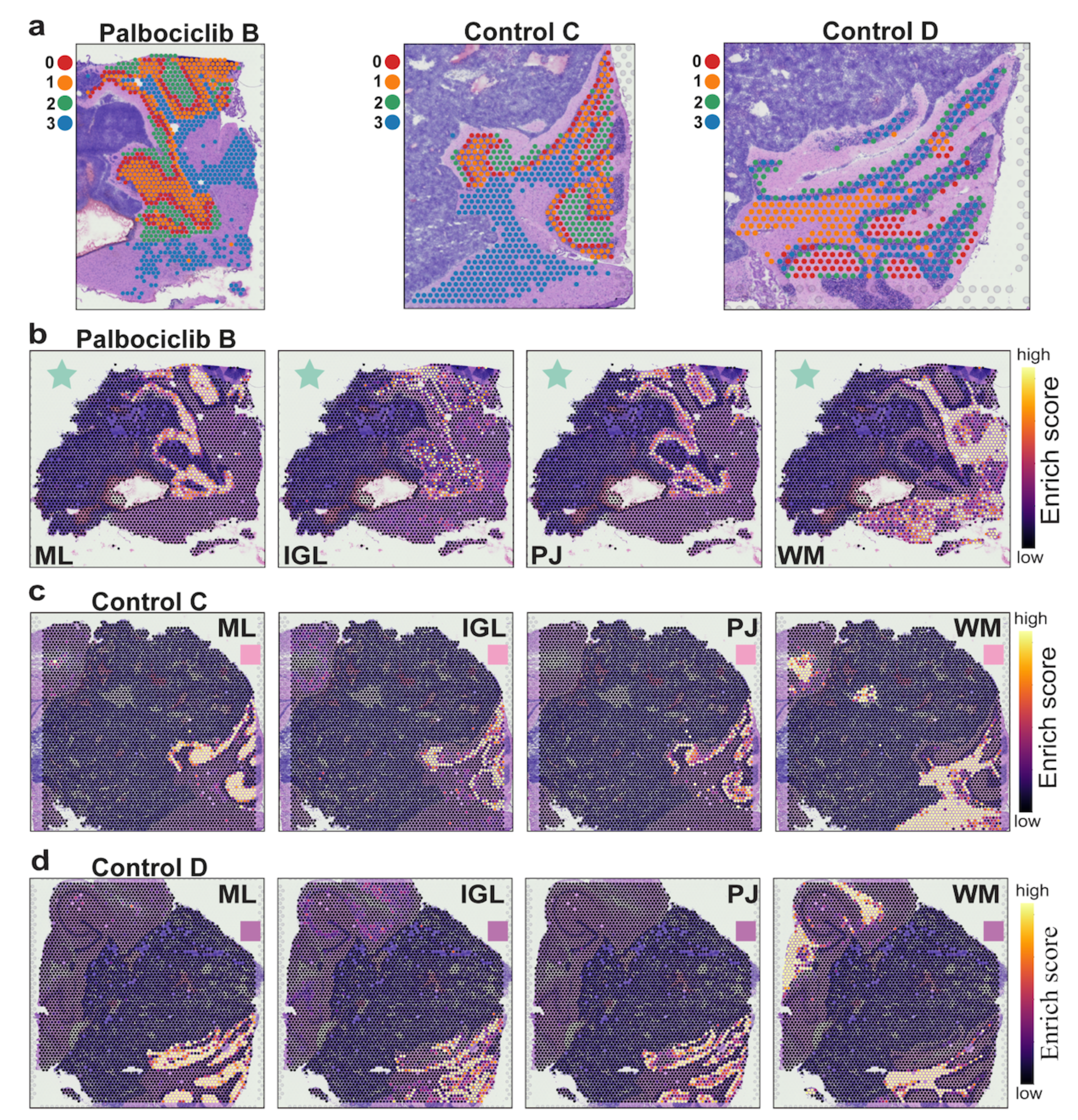
Supplementary Figure S2: Structures of mouse histology can be identified using unsupervised clustering across each Visium spatial transcriptomics samples, with high resolution and additional information about heterogeneity in gene expression even within one cluster. A-C. Unsupervised clustering results for mouse tissue regions for samples Palbociclib B, Control C, and Control D, respectively; showing the clustering strongly corresponds to known cerebellum structures clear from the H&E. Colours and numbers indicate different clusters. B-D. Per-spot enrichment results for top 10 marker genes of each cluster. Data for samples Palbociclib A for sample Palbociclib B, Control C, and Control D are shown. Regions of high enrichment correspond to the clustering results and well-defined cerebellum structures (ML: Molecular layer, PJ: Purkinje, IGL: internal granule layer, WM: White Matter layer).

**Supplementary Figure S3.**
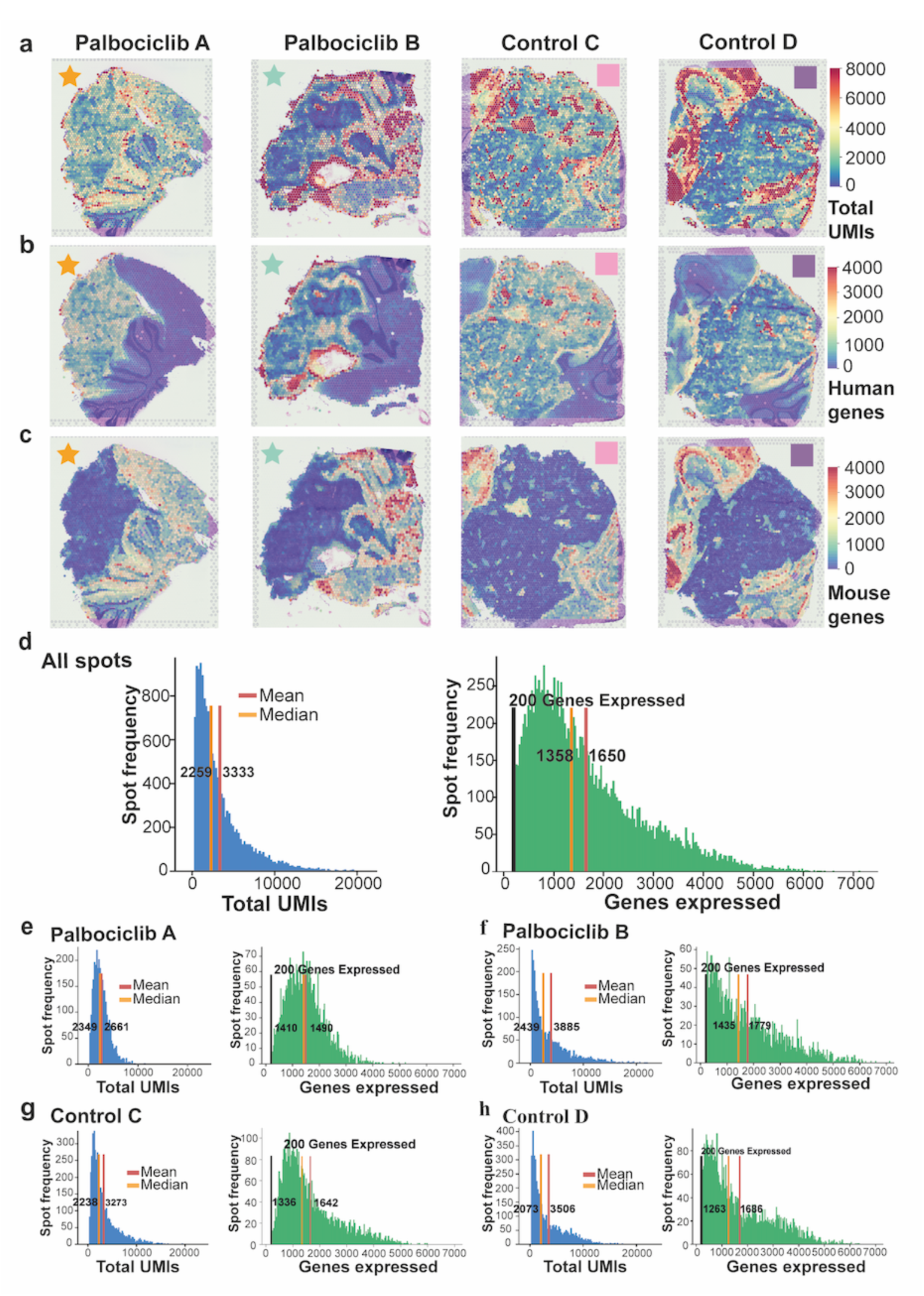
Supplementary Figure S3. Spatial visualisation and quality control assessment of gene expression within Medulloblastoma PDO × (MB-PDOX) mouse brains. A. The total number of unique molecular identifiers (UMIs) per spot are visualised at their position within MB-PDX sections. The spots are coloured according to their detected UMI content (blue - low number of UMIs, yellow = medium number of UMIs, red = high number of UMIs). B. Visium spatial RNA-seq detects human genes within the human tumour regions. The numbers of human genes detected for each spot across for each sample are shown (blue = low number of human genes, yellow = medium number of human genes, red = high number of human genes). C. Equivalent to B, but for mouse genes only. D. Histograms show the distribution of total UMIs detected for data from all spots across all samples (left, blue) and that for the number of genes expressed (right, green). The mean and median of each respective measurement are indicated in-text next to red and orange vertical lines, respectively. The black vertical line on the green histogram indicates a minimum of 200 genes expressed per spot was used to filter extremely low-quality spots. E-H. The same as D, except for data is for each separate sample.

**Supplementary Figure S4.**
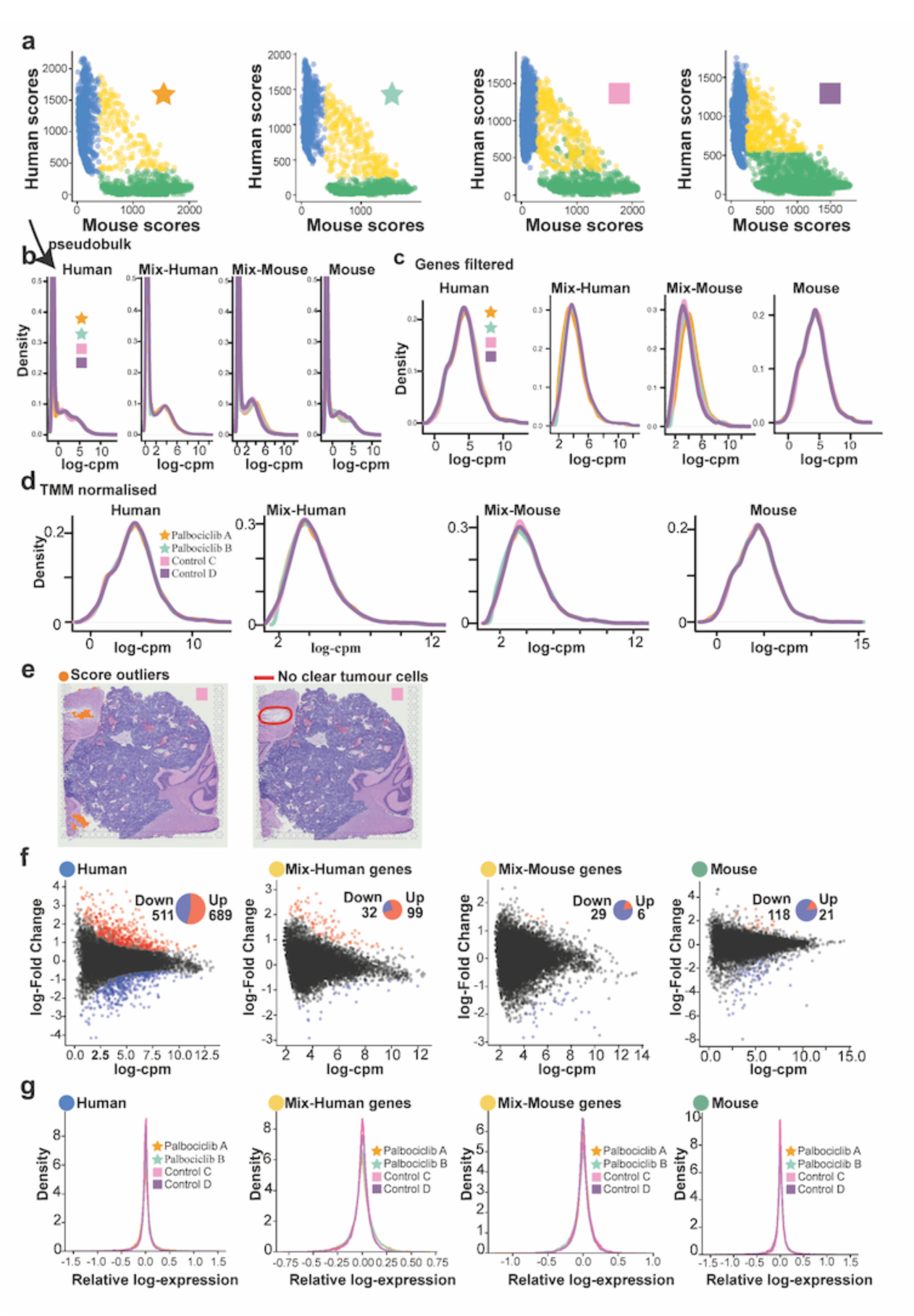
Supplementary Figure S4. Diagnostic plots to assess technical variation and normalisation for gene expression analysis of Visium ST-seq data. Scatter plots with the sum of normalised human gene expression on the y-axis (normalised UMI count) and mouse gene expression on the x-axis per spot, allowing for classification of spots into species type. Spots are classified as human (green), mix (yellow, containing both human and mouse genes) or mouse (blue) based on whether they predominantly express one species' genes or a combination of both (see Methods). The distributions of gene expression as raw log-cpm (log-counts-per-million) for each sample after pseudobulking (see Methods) for each of the human (left), mix (middle; including human genes in mix or mouse genes in mix, denoted as mix-human and mix-mouse respectively) and mouse (right) spots. C.The same data as B, except that data were filtered to remove lowly expressed gene based on log-cpm cutoffs. The very high peaks for lowly expressed genes in B are not observed and the distributions become closer to Gaussian distributions. D.The same data as B, except for log-cpm normalisation using trimmed-mean-squares (TMM) to estimate size factors after gene filtering. After filtering the data were more normally distributed. No sign of technical variation was observed among the four samples. E. Adjustment of spot assignment based on pathological annotation. (Left) outlier spots in sample Control C which were relabelled as'mouse'as opposed to'mix'spots (see Methods). (Right) pathologist annotation indicating no clear evidence of tumours when examining the histology which motivated the relabelling of the outlier spots. F. Scatter plots showing differentially expressed genes detected across the range of gene expression, suggesting that data were appropriately normalised with no bias towards highly or lowly abundant genes. Each point in the scatter plot is a gene, with red points indicating upregulated genes under Palbociclib treatment in MB and blue indicating downregulated genes.The y-axis is the Iog2-Fold Change from the Limma-Voom differential expression (DE) analysis comparing Palbociclib-treated against control samples. The x-axis is the Iog2-cpm average gene expression. The pie charts display the proportion of up-and down-regulated genes. From left-to-right, the described scatter plot is repeated for DE analyses comparing pseudobulked human spots/genes, mix spots with human genes, mix spots with mouse genes, and mouse spots/genes. G. The density of the Relative Log Expression (RLE) values for each gene in each sample pseudobulked (from left-to-right) by human spots/genes, mix spots/human genes, mix spots/mouse genes, and mouse spots/genes. A distribution around 0 for each sample indicates the data were appropriately normalised for differential expression analysis with no systematic shift in the relative gene expression between the pseudobulked samples in each comparison.

**Supplementary Figure S5.**
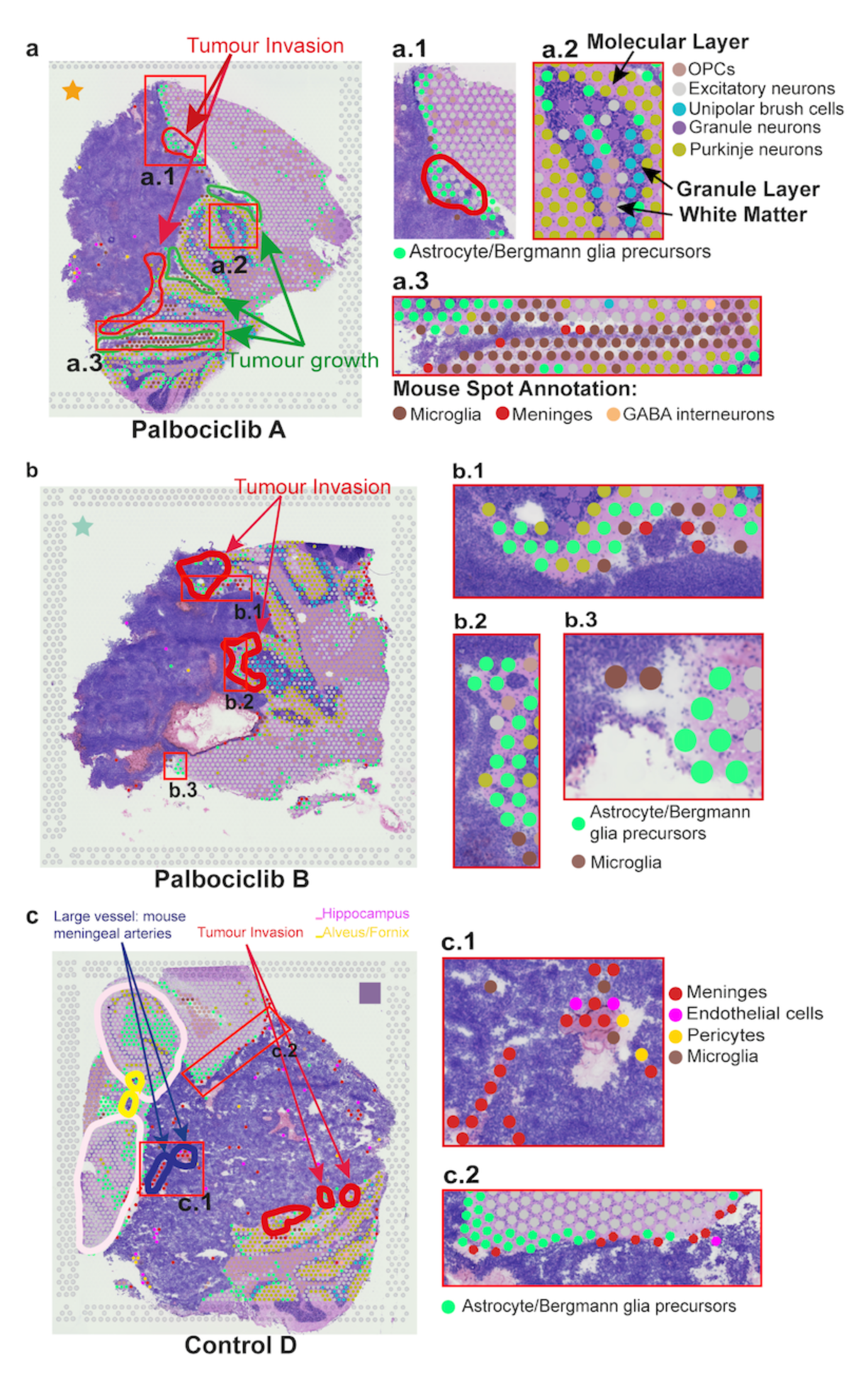
Data-driven cell type detection for spatial transcriptomics spots highly correlated with pathologist's annotation based on mouse histology and had a high resolution sufficiently reflecting cell-type heterogeneity within small tissue regions. A. Independent pathology annotation (shown as contours/circles) of sample Palbociclib A overlaid on top of Visium spots cell types (shown as different colours). A1-A3. Enlarged regions in Palbociclib A (red boxes). A1. Enlarged region showing astrocytes at the tumour/mouse interface, and a tumour invasion region annotated by the pathologist in red. A2. Enlarged region showing the different layers of the mouse cerebellum and the spot cell types; indicating the expected cell types in their respective cerebellum layers. Purkinje cells were primarily in the molecular layer; granule, unipolar brush, and excitatory neurons were dominant in the granule layer; while the white matter was predominantly oligodendrocyte precursor cells (OPCs). A3. Enlarged region annotated by the pathologist as 'tumour growth'; the dominant mouse cell types interfacing with the tumour are microglia, astrocyte/Bergmann glia, and meninges.B. Independent pathology annotation of sample Palbociclib B overlaid on top of Visium spots cell types. B1-B3. Enlarged region in Palbociclib B (red boxes). B1-B2. Enlarged interface region from B (red boxes) which overlaps pathologist annotation 'Tumour invasion' indicates astrocytes, microglia, and meninges as dominant cell types. B3. Enlarged interface region indicates astrocytes and microglia as the dominant cell types in the interface spots. C. The same as A, B for sample Control D. C1-C2. Enlarged region in Control D (red boxes). C1. Enlarged region annotated by the pathologist as Large vessel: mouse meningeal arteries'. Meninges, endothelial cells, pericytes, and microglia are the predominant cell types in these regions. The dominant cell type for these areas are primarily blood-vessel related, in concordance with the pathologist annotation. C2 Enlarged region from sample Control D showing the tumour-mouse interface, astrocytes, meninges, and

**Supplementary Figure S6.**
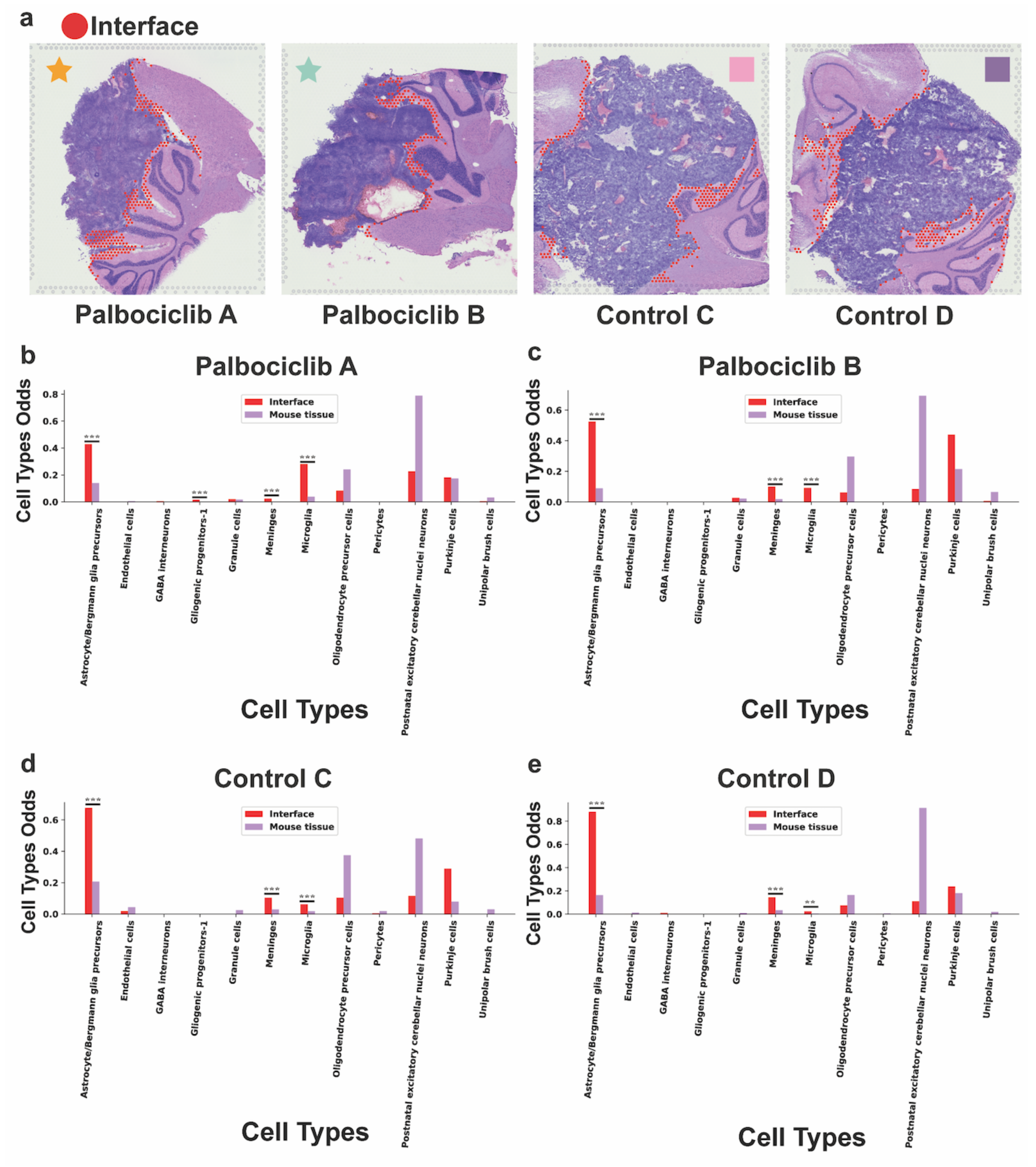
Definition of spots at the tumour-mouse interface. A. Spots at the tumour-mouse interface which were used to test for enrichment of astrocytes for samples (left-to-right) Palbociclib A, Palbociclib B, Control C, and Control D, respectively. B-E. Gene expression of each cell types between interface and "mouse tissue" for samples Palbociclib A, Palbociclib B, Control C, and Control D, respectively.

**Supplementary Figure S7.**
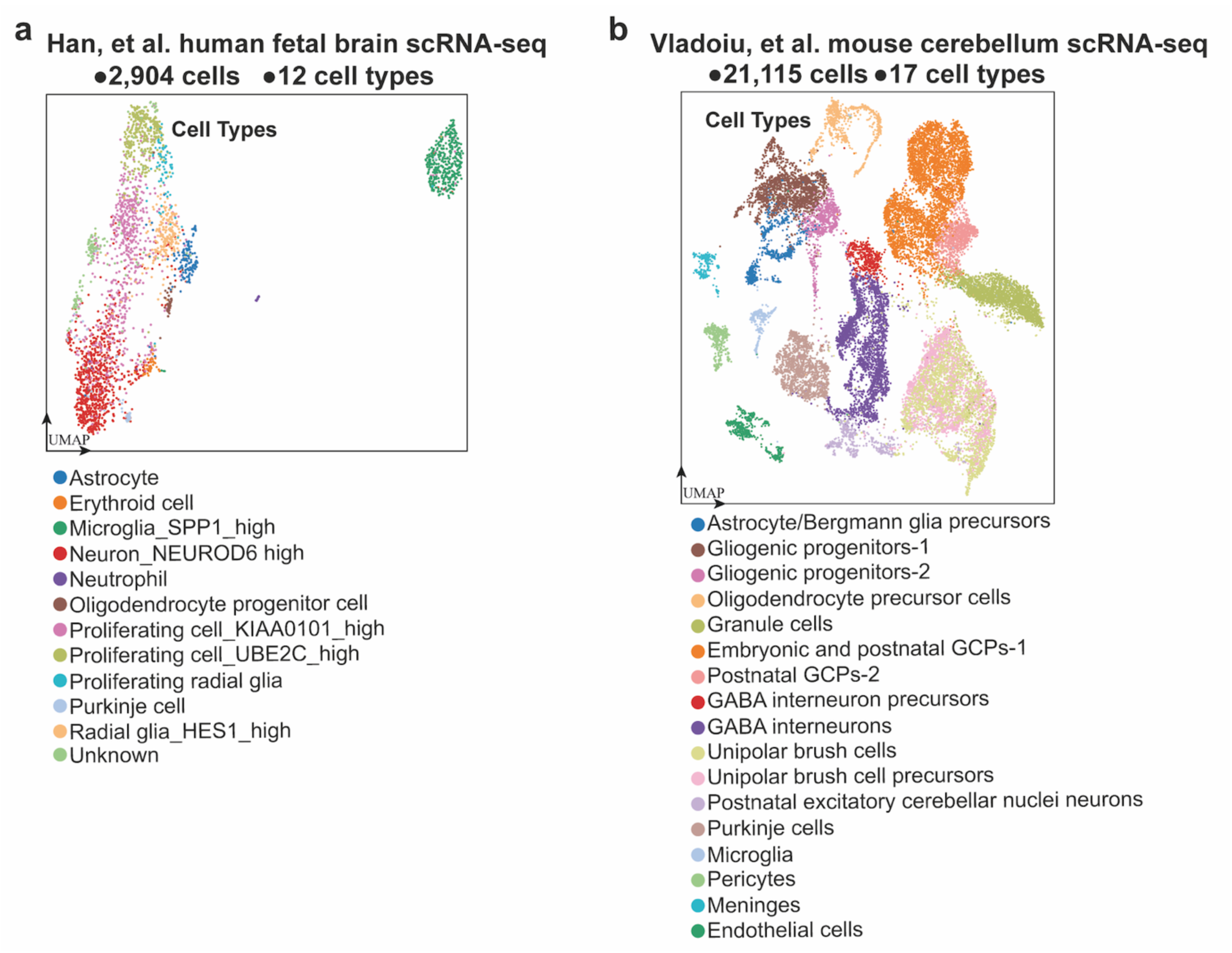
Overview of the reference single cell RNA-seq data used for automated spot cell type identification. A. The reference scRNA-seq data used to annotate human/mix spots (only human genes were used for the mix spots). The data was obtained from Han, et al. (2020), consisting of 2,904 cells spanning 12 different cell types from 13 week male human foetal brain. B. The reference scRNA-seq data used to annotate mouse/mix spots using mouse genes. The data was obtained from Vladoiu, et al. (2019), consisting of 21,115 cells spanning 17 cell types once subsetted to cell types only present at postnatal day 7 (P7).

**Supplementary Figure S8.**
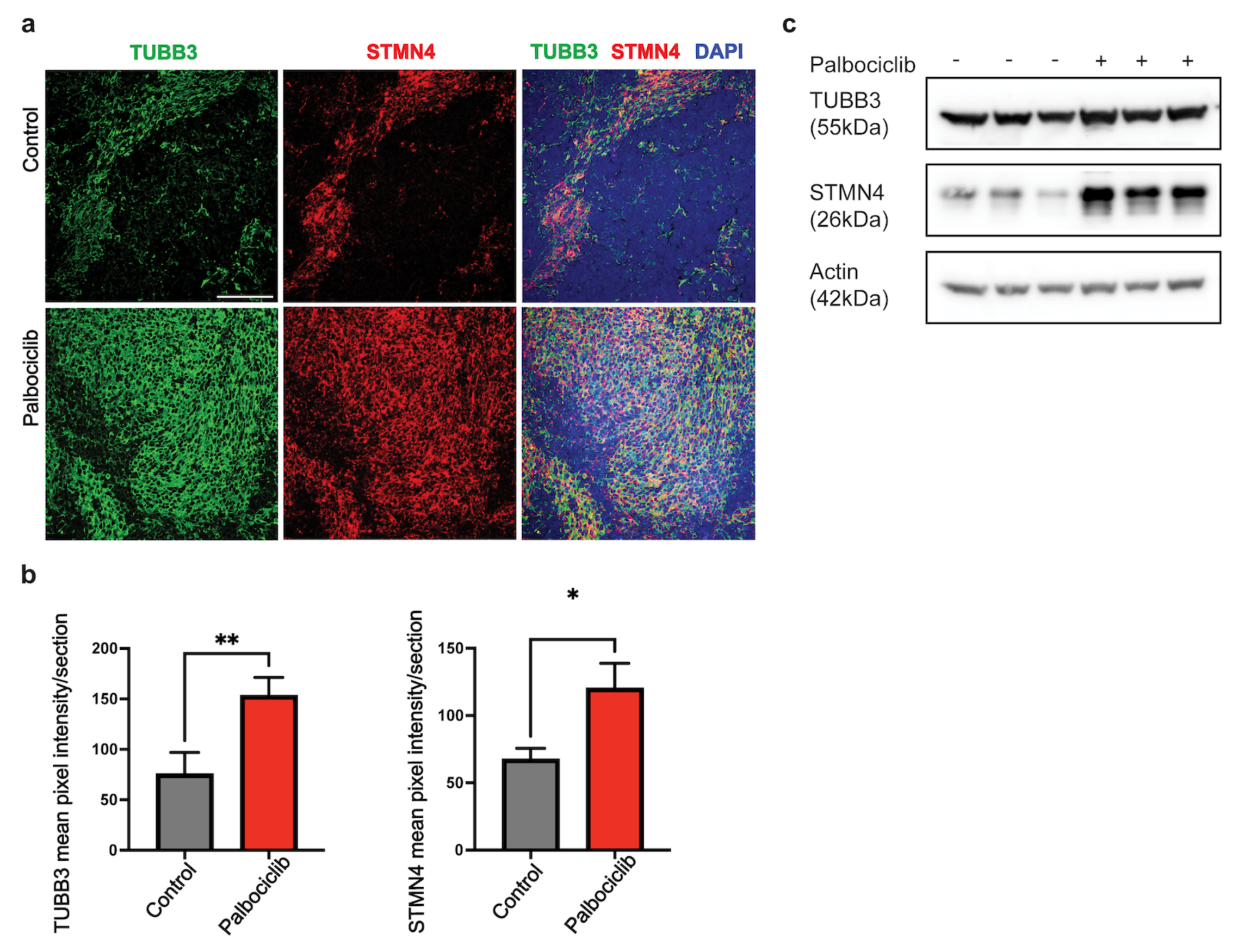
Palbociclib-treatment induces neuron differentiation in SHH MB. Representative images of two markers of neural differentiation (TUBB3 and STMN4) in immunofluorescence-labelled Med-1712FH orthotopic tumours following Palbociclib treatment (a, top panel) compared to untreated control tumours (a, bottom panel). Sections were counterstained with DAPI. Scale bar 100 μm. Quantitative analysis of TUBB3 (b, left) and STMN4 (b, right) staining in Med-1712FH tumours. Mean pixel intensity was quantified for each marker for vehicle (n = 3) and drug-treated tumours (n = 3) using ImageJ software. Data are presented as the mean ± SEM. Statistical evaluation was performed using an unpaired two-tailed t-test (? = 0.05) with Welch's correction. Statistically significant differences are indicated (*p < 0.05; **p < 0.01). (c) Immunoblot of protein extracts from untreated (n = 3) and Palbociclib-treated (n = 3) Med-1712FH orthotopic tumours for expression ofTUBB3, STMN4 and Actin. Immunoblots were imaged using the BioRad ChemiDoc MP Imager. Supplementary Figure S9. Per-spot enrichment analysis of DE neuron differentiation genes and neuronally differentiated SHH-C subpopulation corroborates pathologist "pale island" annotations.
a. Comparision between pathologist annotations from H&E images (left) and per-spot enrichment scores of differentially expressed (DE) neuron differentiation genes and SHH-C in Palbociclib A. 'Pale island' annotations are indicated in yellow, with enlarged sections in red boxes, b-c. Equivalent to A except for samples Control C and Control D.

**Supplementary Figure S9.**
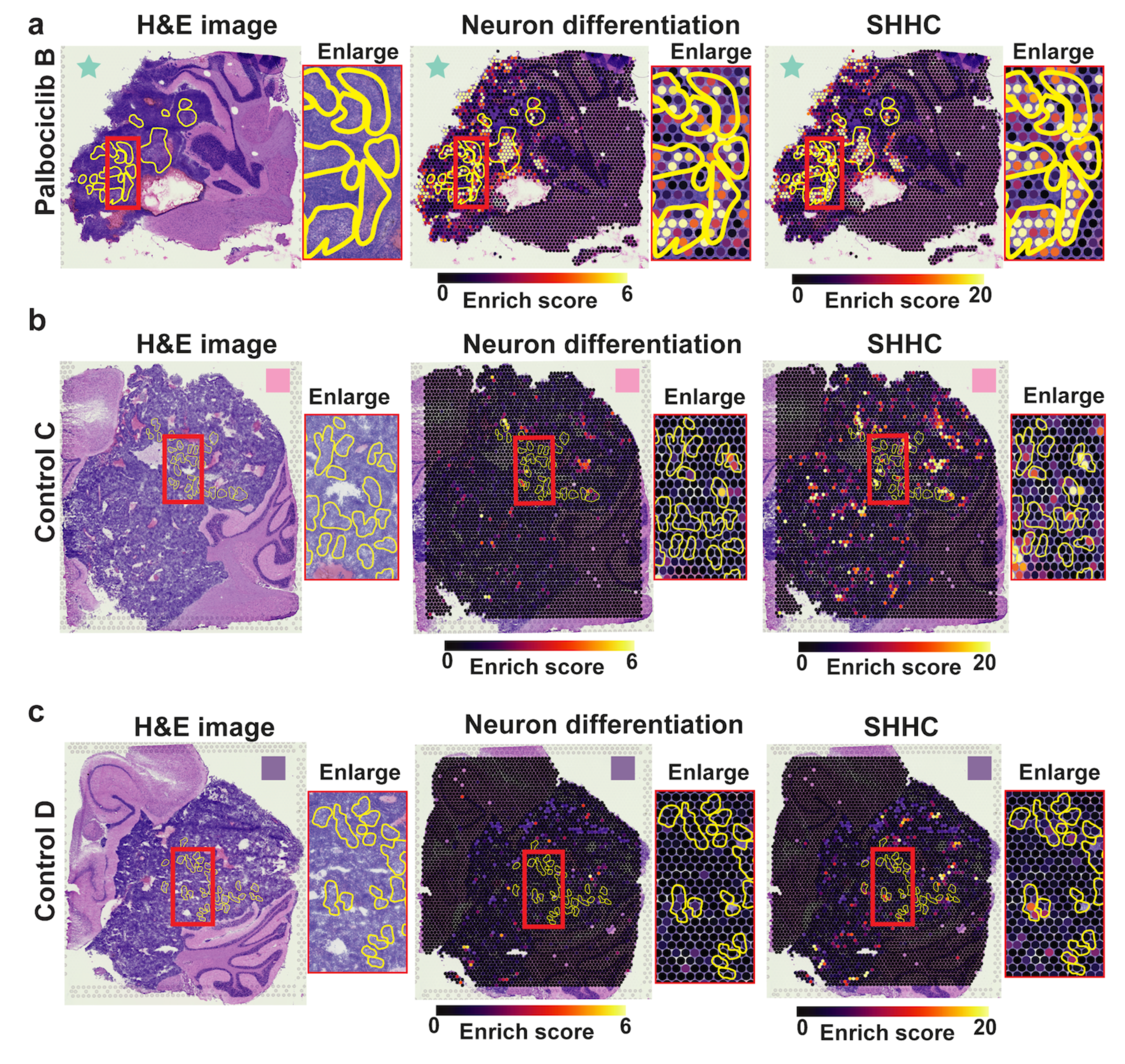
Per-spot enrichment analysis of DE neuron differentiation genes and neuronally differentiated SHH-C subpopulation corroborates pathologist "pale island" annotations. **a.** Comparision between pathologist annotations from H&E images (left) and per-spot enrichment scores of differentially expressed (DE) neuron differentiation genes and SHH-C in Palbociclib A. 'Pale island' annotations are indicated in yellow, with enlarged sections in red boxes, **b-c.** Equivalent to A except for samples Control C and Control D.

**Supplementary Figure S10.**
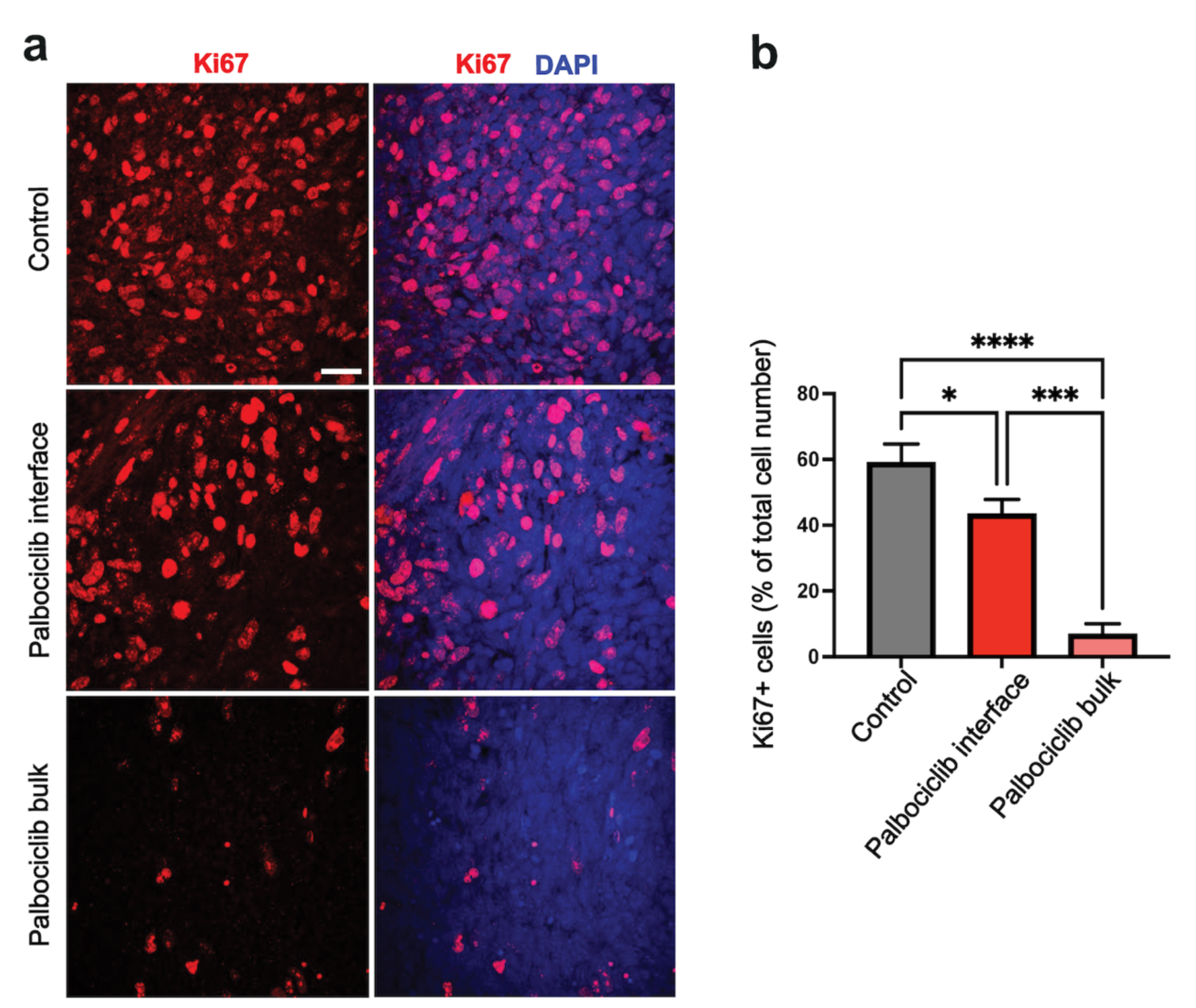
Spatial response of cell proliferation to Palbociclib treatment in SHH MB. Representative images of cell proliferation (K?67) in immunofluorescence-labelled Med-1712FH orthotopic tumours in untreated control tumours (a, top panel) compared to the interface (a, middle panel) and bulk (a, bottom panel) regions of tumours following Palbociclib treatment. Sections were counterstained with DAPI to determine total cell number. Scale bar 20 μm. Quantitative analysis of tumour cells staining positive for K?67 (b) in untreated control tumours (n=3) compared to the interface and bulk tumour regions of Palbociclib-treated (n=3) tumours. Data are presented as the mean ± SEM. Statistical evaluation was performed using a multiple t-test with Tukey's correction. Statistically significant differences are indicated (*p < 0.05; ***p < 0.001 ? ****p < 0.0001).

**Supplementary Figure S11.**
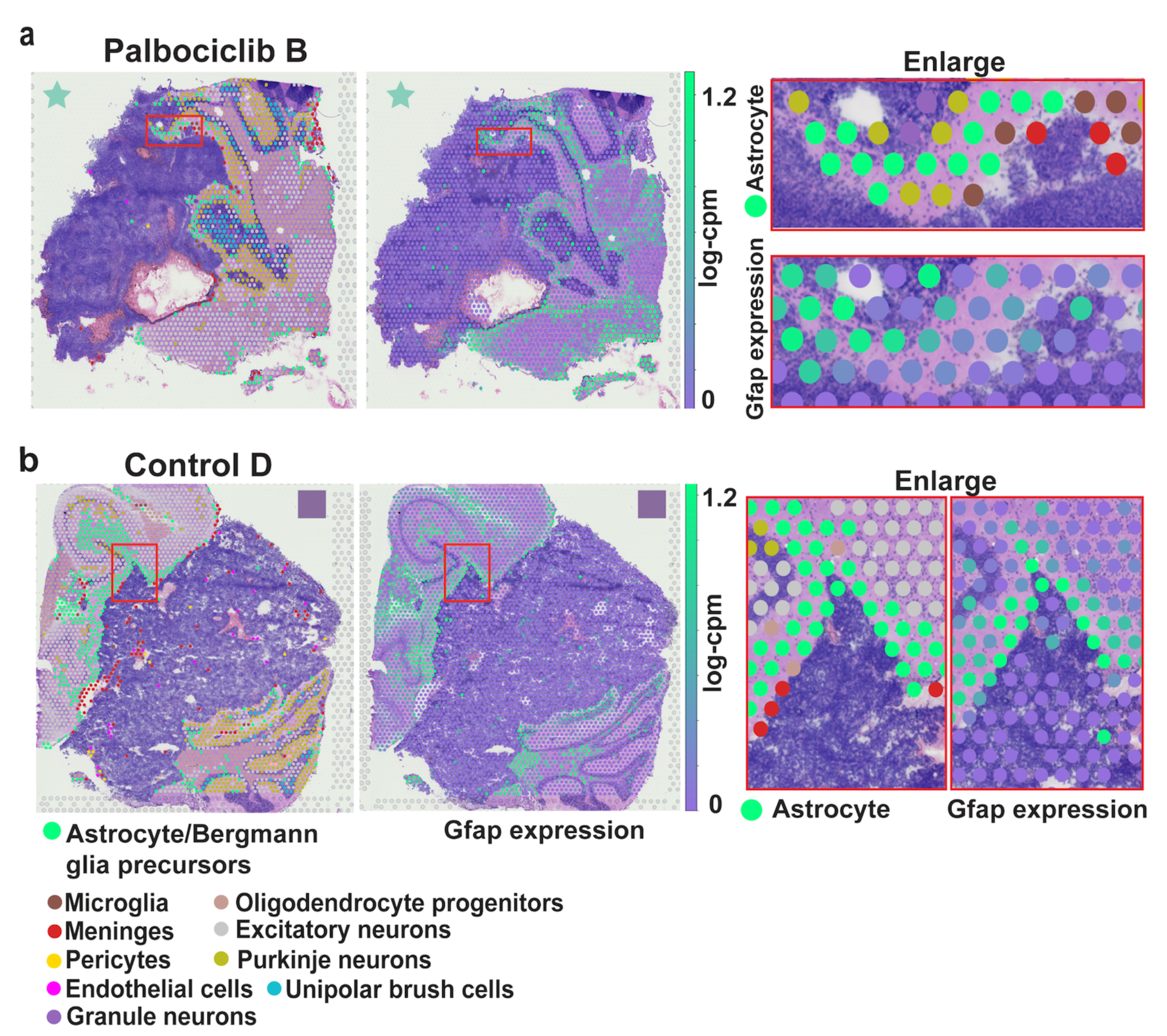
Analyses of mouse-only tissue region shows that mouse astrocytes localise to the tumour-mouse interface. **a.** Sample Palbociclib A with overlaid mouse spot annotations (left) and Gfap gene expression (right, in log-cpm (log-counts-per-million)). The red box indicates the region of interest enlarged to the far right. Cell types are displayed as different colours. Overall, there is concordance between Gfap gene expression in spots and astrocyte cell types; with both localised to the interface region, **b.** Equivalent to **a**, except depicting sample Control D.

**Supplementary Figure S12.**
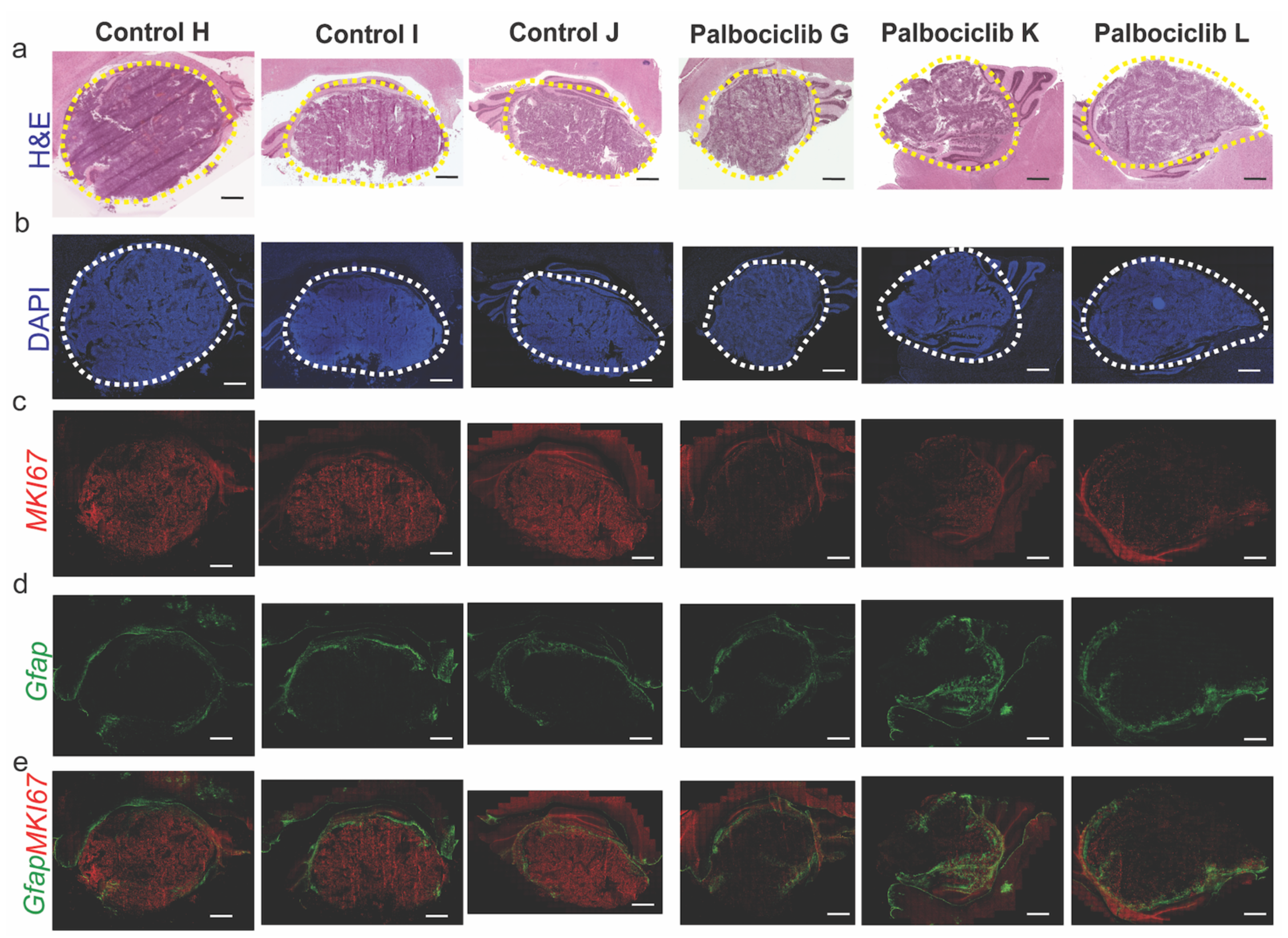
The expression of cell proliferation marker MKI67 is localised tumour regions with expression of astrocyte marker, Gfap. (a) High resolution H&E images of drug-treated and untreated PDOX with tumour volumes outlined in yellow dashed lines, (b) DAPI staining of control and drug-treated PDOX with tumour volumes outlined in white dashed lines, (c) Target RNA molecule expression of cell proliferation marker MKI67 using RNAscope in control and drug-treated PDOX. (d) Target RNA molecule expression of astrocyte marker Gfap localised to the tumour-mouse interface region, (e) Merged image of MKI67 expression with Gfap expression across drug-treated and untreated PDOX.

**Supplementary Figure S13.**
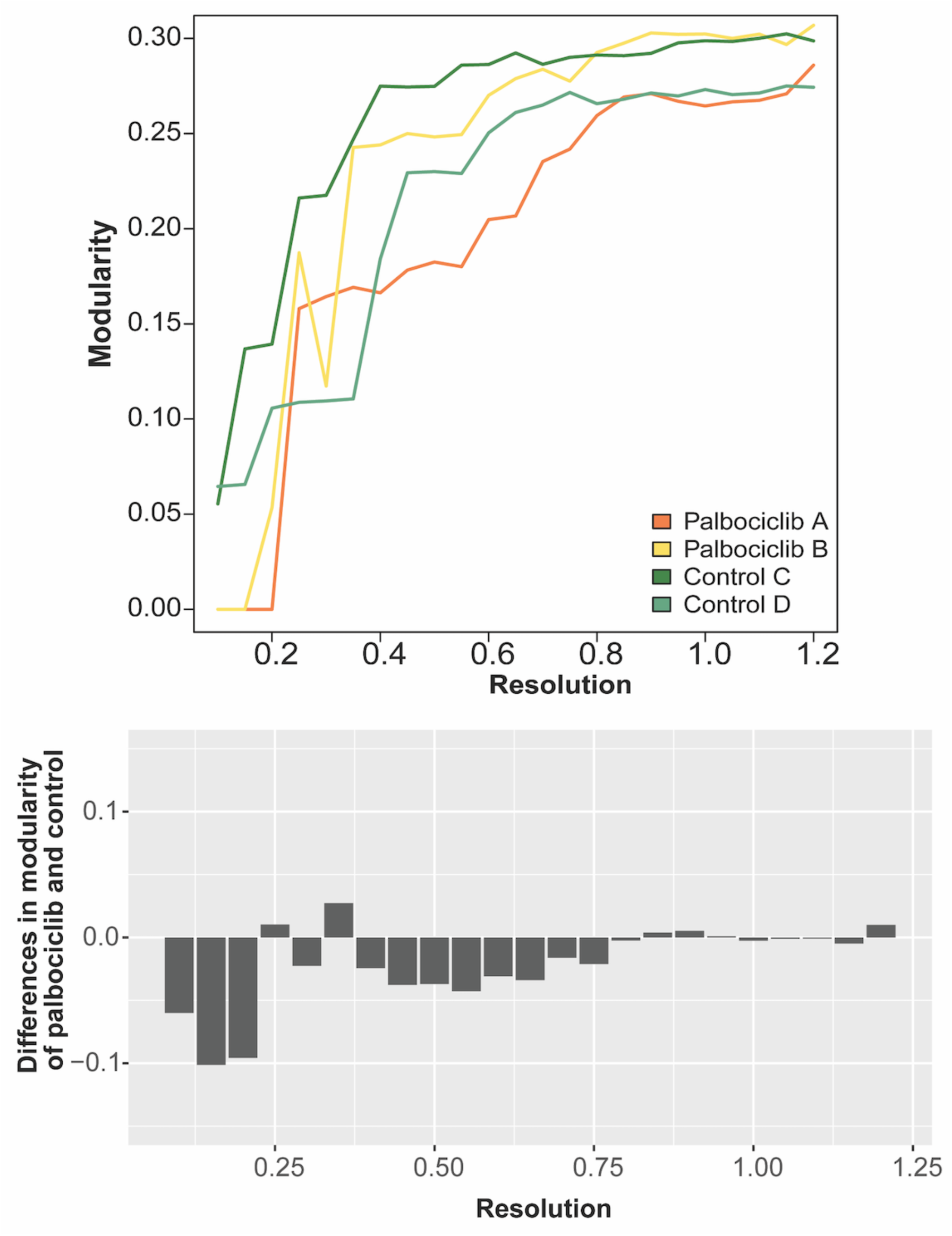
Modularity per sample estimated from each clustering resolution used in Figure 3a. The x-axis is the same clustering resolution in panel A. The y-axis is the Shannon entropy per sample. Colours represent different individuals. The bar plot on the bottom panel shows the differences in Shannon entropy of Palbociclib and control (i.e. mean of entropy in Palbociclib - mean of entropy in control).

